# Replication stress alters CENP-A nucleosome stability during S phase

**DOI:** 10.1101/2025.05.23.655122

**Authors:** Alexander S. Lee, Che-Fan Huang, Taojunfeng Su, Justin Bodner, Neil L. Kelleher, Daniel R. Foltz

## Abstract

The maintenance of centromere identity is essential for the proper segregation of chromosomes during cell division. Centromere identity is epigenetically specified by centromeric histone H3 variant CENP-A, and its retention during DNA replication is facilitated by HJURP chaperone. Replication stress disrupts replication fork progression and can negatively influence the interactions between histone chaperone network necessary for retention and deposition of parental and new histones, respectively. In this study we investigate the role of replication stress response on centromere inheritance. We define changes in centromere-associated proteins that govern stability of centromeric and canonical nucleosomes through proximity labeling coupled with affinity purification mass spectrometry. We identified that under replication stress, CENP-A-containing chromatin strongly enriches for SWI/SNF chromatin remodeling proteins ATRX. We show that depletion of ATRX and its associated histone H3.3 chaperone DAXX results in the loss CENP-A retention in S-phase and loss persists into the subsequent cell cycle. Altogether our findings provide insight into how replication stress negatively influences centromeric chromatin instability and delineates a function of DAXX-ATRX complex in maintaining centromere inheritance during DNA replication.

## 2. Introduction

Human centromeres are essential for proper chromosome segregation and are often sites of genomic instability (1). The underlying centromeric chromatin are specialized chromatin domains that facilitate recruitment of constitutive centromere-associated network (CCAN) components and kinetochore assembly for proper chromosome segregation (2–5). Centromere protein A (CENP-A) is a histone H3 variant that epigenetically specifies centromeric chromatin (6). *De novo* of deposition CENP-A occurs in early G1 phase and is facilitated by CENP-A deposition pathway involving its dedicated histone chaperone HJURP, and Mis18 complex (6–10).

Human centromeric DNA is replicated during mid to late S-phase, and the retention of parental CENP-A is critical for re-establishing centromeric chromatin, as its expression and deposition are not coupled to DNA replication compared to canonical histone H3.1 (11–13). The retention of parental CENP-A is facilitated by HJURP in coordination with MCM2-7 complex, in similar manner to ASF1-MCM2-7 complex for retention of parental H3.1 (14–18). During the replication of centromeric chromatin, H3.1 and H3.3 is deposited in regions unoccupied by parental CENP-A and are thought to serve as placeholders for deposition of new CENP-A in G1 (19). Thus, it is critical that parental CENP-A are retained efficiently as to prevent the loss of centromere identity.

Human centromeres are rich in alpha-satellite DNA repeats which are organized in a head-to-tail orientation of AT-rich 171 bp monomers forming higher-order repeats arrays (HOR). The repetitive nature of HORs makes these regions more prone to DNA recombination events, and forming higher order DNA structures including non-B DNA conformations and DNA:RNA R loops, thus making it a challenge to replicate centromere regions (1,20–23). These nucleic acid structures can be endogenous sources of replication stress, which is characterized by the slowing or stalling of replication forks (24,25). Specialized replication and repair mechanisms involving translesion synthesis polymerases, topoisomerase TOP2A, and DNA loop extrusion SMC5-6 complex have been proposed to prevent centromere defects in the context of genomic instability (26). Similarly, the H3.3 DAXX chaperone has been implicated in centromere stability through independent mechanisms involving the prevention of R loops and its restriction to PML bodies (27,28). Although previous studies have demonstrated mechanisms to prevent centromere maintenance in the context of genomic instability, the link between replication stress and epigenetic inheritance of centromeres is not well understood.

In general chromatin, the retention of parental and deposition of new canonical histones are tightly coupled with DNA replication (17,29–34). As such, the fidelity of canonical histones can be negatively influenced in the presence of replication stress. Replication stress can lead changes in histone modifications and erroneous re-distribution of both parental and newly synthesized histones by shunting them to various histone chaperone deposition pathways (35–38). Thus, the erroneous re-distribution of both parental and newly synthesized histones is thought to lead to changes of chromatin states (37,39).

Compared to general chromatin, the relationship between replication stress and centromeric chromatin inheritance is less understood. Rapid CENP-A depletion during DNA replication can lead to the formation of R-loops at centromeres and is associated with increased incidences of chromosome translocation events, anaphase bridges, and chromosome breakages – all of which are characteristics of chromosomal instability (23). In addition, the rapid depletion of HJURP during S-phase leads to loss of CENP-A (14). The stable inheritance of CENP-A suggests a protective function in centromere stability, and that loss of CENP-A contributes to downstream replication stress. We investigate how replication stress influences the CENP-A retention at centromeres and delineate the protein-protein interactions that contribute to centromere instability. We observed centromeric loss of CENP-A in the presence of acute replication stress caused by various genotoxic agents. Coupling both *in vitro* proximity labeling and biotin purification mass spectrometry, we defined the protein interaction network at both centromeric and general chromatin during DNA replication and replication stress. We identified an association between CENP-A and the SWI/SNF chromatin remodeling protein ATRX with centromeric chromatin that is enhanced upon replication stress, and the loss of ATRX or its H3.3 chaperone DAXX results in loss of CENP-A during centromere replication.

## 3. Results

### Acute DNA replication stress leads to reduced parental CENP-A retention in S-phase

While loss of CENP-A contributes to replication stress and genomic instability at centromeres (23,40), how replication stress influences the fidelity of CENP-A retention is unknown. Based on the effects of replication stress on histone PTM inheritance, we hypothesized that replication stress leads to loss of CENP-A nucleosomes and centromere identity. To test this, we synchronized non-transformed hTERT-RPE1 (RPE1) cells at G1 phase by CDK4 inhibition by Palbociclib (Palb) treatment, and subsequently released into S-phase in the presence of various agents known to cause replication stress including inhibition of replicative DNA polymerases α and δ by aphidicolin (Aph) **(Figure 1A and B, SI Figure 1)** (41,42). We observed loss of CENP-A at centromeres marked by CENP-T foci when treated with Aph **(Figure 1C)**. Similarly, we observe loss of CENP-A at centromeres by inhibition of nucleotide supply by hydroxyurea (HU) – an inhibitor of ribonucleotide reductase and known replication stress inducing agent **(Figure 1C)** (42–44). To confirm the presence of replication stress, we measured the presence of DNA damage and replication stress marker γH2AX **(Figure 1D).** Thus, these results demonstrate that CENP-A loss in the presence of different inducers of replication stress that target independent mechanisms is a non-specific phenomenon.

**Figure 1:**
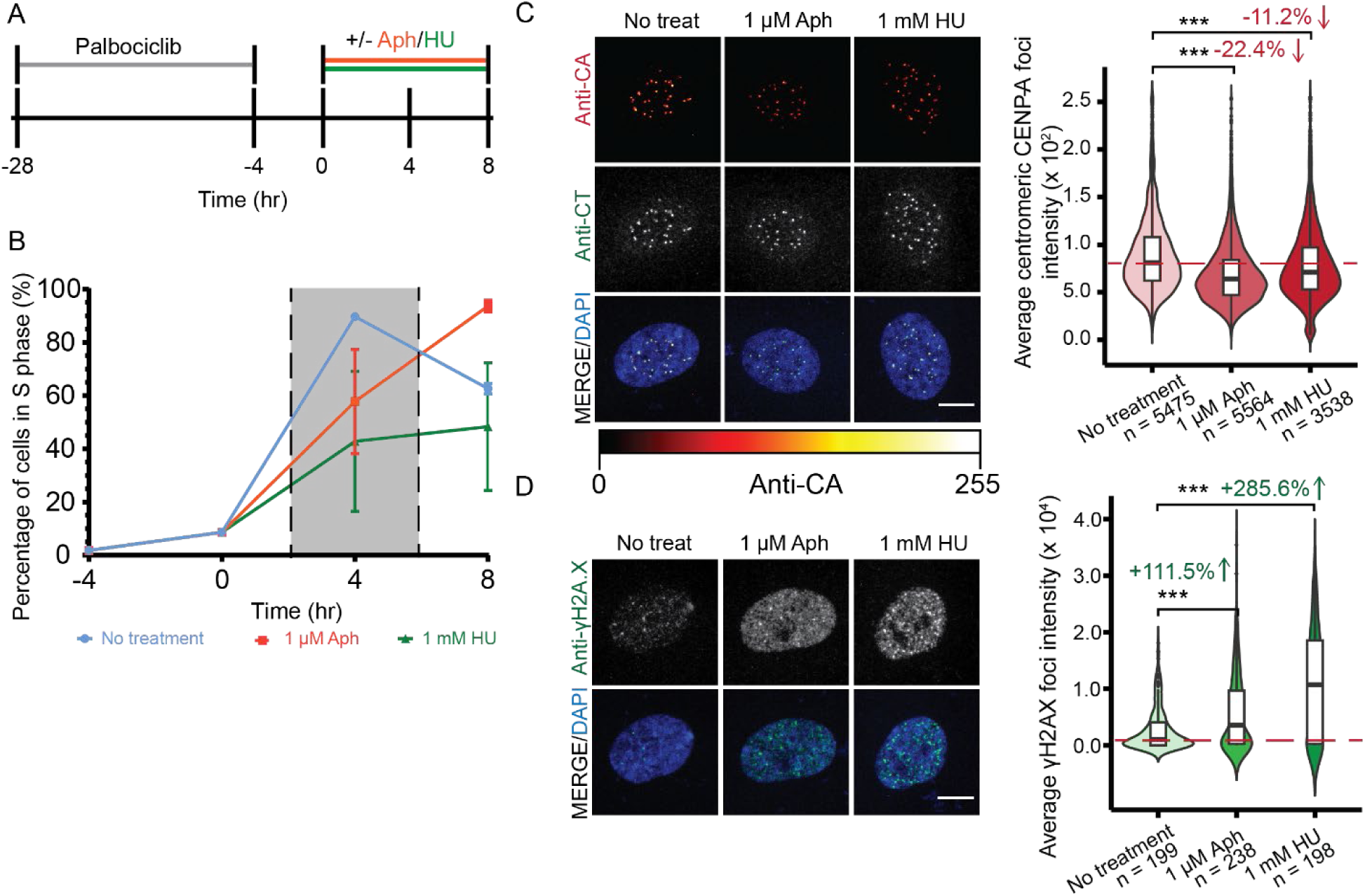
Stress leads to loss of CENP-A at centromeres during DNA replication. **A)** Schematic of experimental design. **B)** Analysis of total cell population migrating through S-phase after post washout of palbociclib with or without replication stress inducing agents. The shaded region represents time-frame of normal S phase progression. **C)** Representative images of RPE1 cells treated with or without replication stress agents (left) and quantification of CENP-A foci co-localized at CENP-T foci (right). n = representing the number of CENP-A foci measured. CENP-A alone is presented as an intensity gradient. Merged images represent CENP-A as red. **D)** Representative images of RPE1 cells treated with or without replication stress agents (left) and quantification of total γH2A.X levels per cell (right). *n* = represents number of cells measured per group. Red line indicates baseline average intensity relative to no treatment control. Not significant (n.s.) = *p* > 0.05, *p* ≤0.05, *p* ≤0.01 = **, *p* ≤0.001 = *** Experiments were performed with three independent biological replicates. Scale bar is 10 μm.

### TurboID identifies novel interactions between histone H3 variants

The interactions between chromatin-associated proteins (CAPs) and chromatin are often weak and transient, owing to multifunctional roles of CAPs in mediating DNA-centric processes (45,46). To capture both stable and transient CAPs that may contribute to the CENP-A retention, we developed Chromatin-TurboID (ChromaTID), a proximity labeling approach for unique chromatin regions by TurboID and identified the associated proteins with biotin purification mass spectrometry (BP-MS) (47,48). To perform ChromaTID, we generated stable cell lines expressing histone H3 variants (H3.1, H3.3, and CENP-A) with C-terminally fused TurboID biotin ligase under a doxycycline-inducible promoter in RPE1 cells **(Figure 2A and Sl Figure 2).** To identify histone-specific association, we compared proteins enriched by LaminA-TurboID nuclear control under asynchronous conditions **(Figure 2B-D)**. Our BP-MS results from asynchronous cells highlight the specificity of the ChromaTID approach for the identification of CENP-A chromatin-associated proteins. The ASF1, NASP, and FACT chaperones are well-established histone chaperones common to H3.1 and H3.3 are associated with both histones. However, absent from CENP-A-TurboID samples, as are the H3.1-specific CHAF1A and CHAF1B, and members of the H3.3 specific CABIN-UBN-HIRA and DAXX-ATRX complexes. H3.3-specific complexes shows some labeling in H3.1-TurboID samples, suggesting these complexes are in proximity **(Figure 2C-E)**. GO term analysis of enriched CAPs shared amongst all three histone H3 variants revealed protein functions associated with DNA metabolic processes including transcription, DNA replication, chromatin remodeling and chromatin organization **(SI Figure 3A-C).**

**Figure 2:**
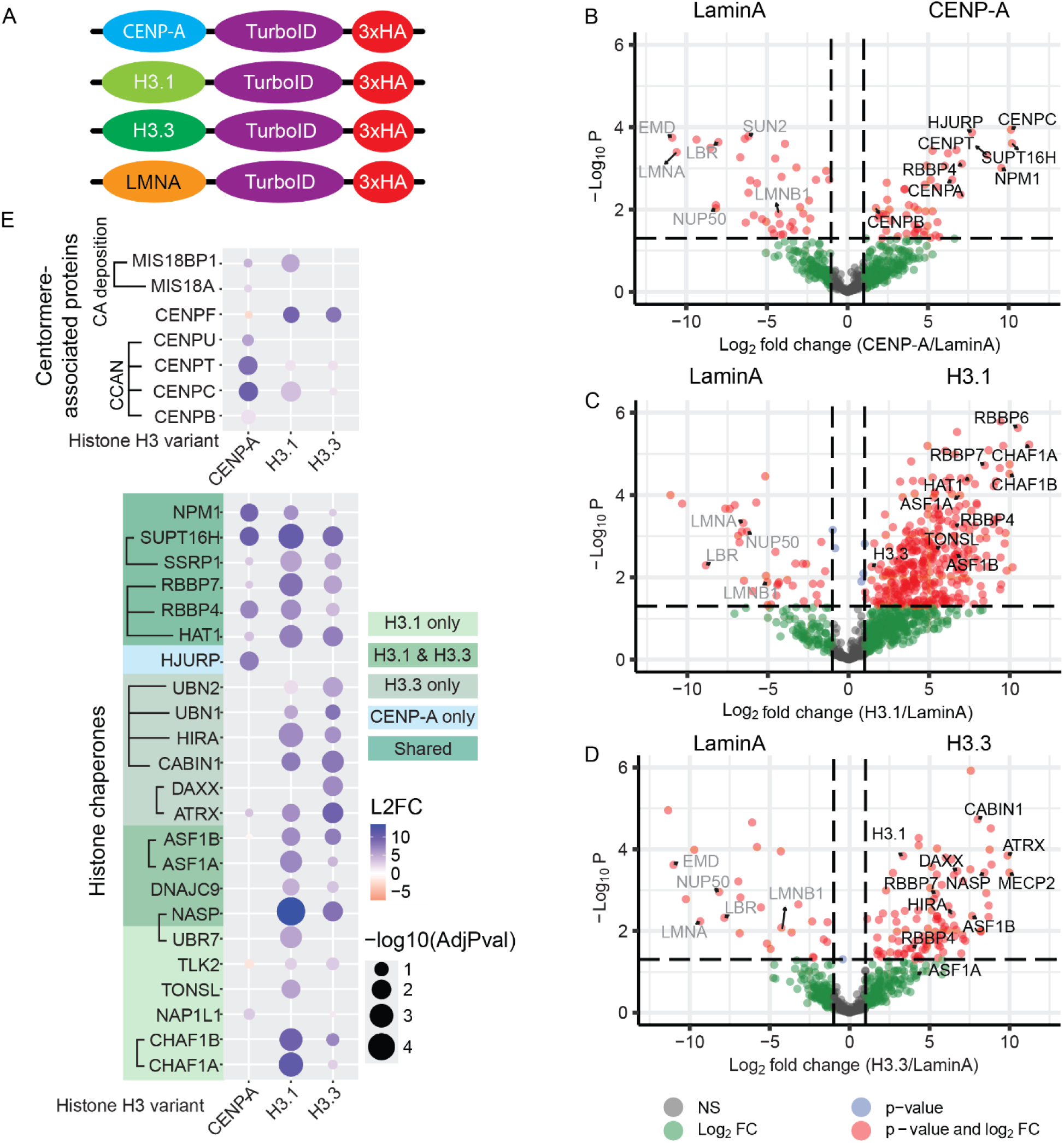
Interaction profiling of replication-coupled and replication-independent histone H3 variants by Chromatin-TurboID and biotin purification mass spectrometry. **A)** Graphical representation of doxcycline-inducible TurboID constructs. **B)** Volcano plots of protein groups identified by BP-MS of randomly cycling CENP-A-TurboID, **C)** H3.1-TurboID, or **D)** H3.3-TurboID cell lines. Three biological replicates were analyzed with three technical replicates per biological replicate. Log2 Fold Change (L2FC) is of [Histone/LaminA control] and plotted against log10 (*p* value). **E)** Bubble plots of statistically significant protein groups identified by BP-MS of randomly cycling TurboID cell lines and classified based on GO-term defined pathways. Three biological replicates were analyzed with three technical replicates per biological replicate. Log2 Fold Change (L2FC) is of [Histone/LaminA control]. Adjusted *p* values were calculated based on bonferroni correction. Brackets indicate known protein complexes.

As expected, CENP-A-TurboID strongly labeled the HJURP chaperone **(Figure 2B and E)**. In addition, our BP-MS results demonstrate enrichment of well-known CENP-A interactions including Mis18 complex and CCAN components **(Figure 2E)** (2,5–7,49). Interestingly, Mis18BP1 is also associated with H3.1-containing chromatin. Likewise, CENP-F, which is a known kinetochore protein is strongly labeled by H3.1 and H3.3-TurboID, consistent with a role in transcription (50–52). Of note, overexpression of CENP-A has been shown to result in specious interactions with H3.3 chaperone HIRA or DAXX and ATRX (53–55). Under our CENP-A-TurboID system and conditions, we did not observe the pronounced interactions with DAXX, ATRX, or HIRA **(Figure 2E)**.

We observed new proximal interactions between CENP-A including interactions with type II topoisomerase TOP2A and TOP2B and chromatin remodeling protein ERCC6 **(SI Figure 3D)**. ERCC6 is well known for its role in transcription-coupled nucleotide excision repair but has demonstrated to participate in histone eviction and DNA double strand break repair (56–58). Unexpectedly, we identified CENP-A interactions with various RNA processing factors including DEAD-box RNA helicases DDX18, DDX50, and RNA deaminase ADAR **(SI Figure 3F)** which may reflect transcriptional activity and R-loop formation at centromeres (59–61). These results demonstrate the specificity of our ChromaTID system to capture histone-specific interactions and differentiate interactions found at centromeric and general chromatin.

### Replication stress leads to global interactome changes in general and centromeric chromatin

To delineate mechanisms that could contribute to the loss of centromeric CENP-A, we investigated CAPs that interact with centromeric chromatin during both normal replication and replication stress. ChromaTID cell lines were synchronized at G1 using Palb, and subsequently released into S phase in the absence or presence of replication stress by Aph treatment **(Figure 3A)**. Our BP-MS results highlight global proteome changes involved in various biological processes including chromatin remodeling, DNA damage response, histone deposition pathways, and centromeres **(Figure 3 and SI Figure 4)**. We identified several new interactions between CENP-A under replicating and stressed conditions including histone modifying enzymes EHMT1 and ASH1L **(SI Figure 4A)** which deposits H3K9me1/2 and H3K36me3, respectively. These results are further supported by previous observations with H3K9me1/2 and H3K36me3 enriched in pericentric heterochromatin (62) and H3.1/3 nucleosomes found within CENP-A-containing chromatin (63–65). Additionally, we identified CENP-A-specific CAPs involved in chromosome structure including SMCHD1 and TOP2 which are known to be involved in DNA repair **(SI Figure 4A)** (66,67). Interestingly, we observed CENP-A interactions with various E3 ubiquitin ligases ARIH2, ZFP91, and TRIP12, which are further enhanced during replication stress **(SI Figure 4B)**. In yeast, ARIH2 and TRIP12 have been shown to degrade unincorporated canonical histones in a DNA-damage dependent manner (68–70). This finding highlights a possible role of protein degradation in removing evicted CENP-A in response to replication stress, and regulate soluble supply of available CENP-A. Within H3.1 interactions, we observed increased interactions with various high mobility groups protein (HMGs) including HMGAN1, HMGA2, and HMG4 in a replication stress-dependent manner **(SI Figure 4A)**. Interactions with these HMGs may reflect a global change in chromatin structure and organization in the context of replication stress (71). We observed two novel interactions with CENP-A involving SWI/SNF chromatin remodeling proteins ATRX and ERCC6 in the context of replication and replication stressed cells **(Figure 3B and SI Figure 4A)**. ATRX is a multifunctional SWI/SNF chromatin remodeling complex where it is involved in establishment and maintenance of heterochromatin (72,73), gene transcription (74–76), replication stress (77–79), and DNA repair (80–82). Additionally, we observed CENP-A interaction with methyl DNA binding protein MECP2 and this interaction is only found during DNA replication **(SI Figure 4A)**. MECP2 facilitates the recruitment of ATRX to heterochromatin and has been implicated to interact with canonical histones (83–85). ERCC6 is best known for its role in mediating transcription-coupled nucleotide excision repair where it is recruited to paused RNA pol II in the presence of DNA damage and serves as an initiation factor for TC-NER complex formation (86,87). Additionally, ERCC6 can be involved in other DNA repair pathways including DSB repair (57,88,89). ERCC6 has been shown to interact with various histone chaperones including NAPL1 and H3 chaperones ASF1A/B and sNASP highlighting a potential role in histone deposition in general chromatin (87,90). Nonetheless, neither ATRX nor ERCC6 have been shown to be involved in centromere inheritance.

**Figure 3:**
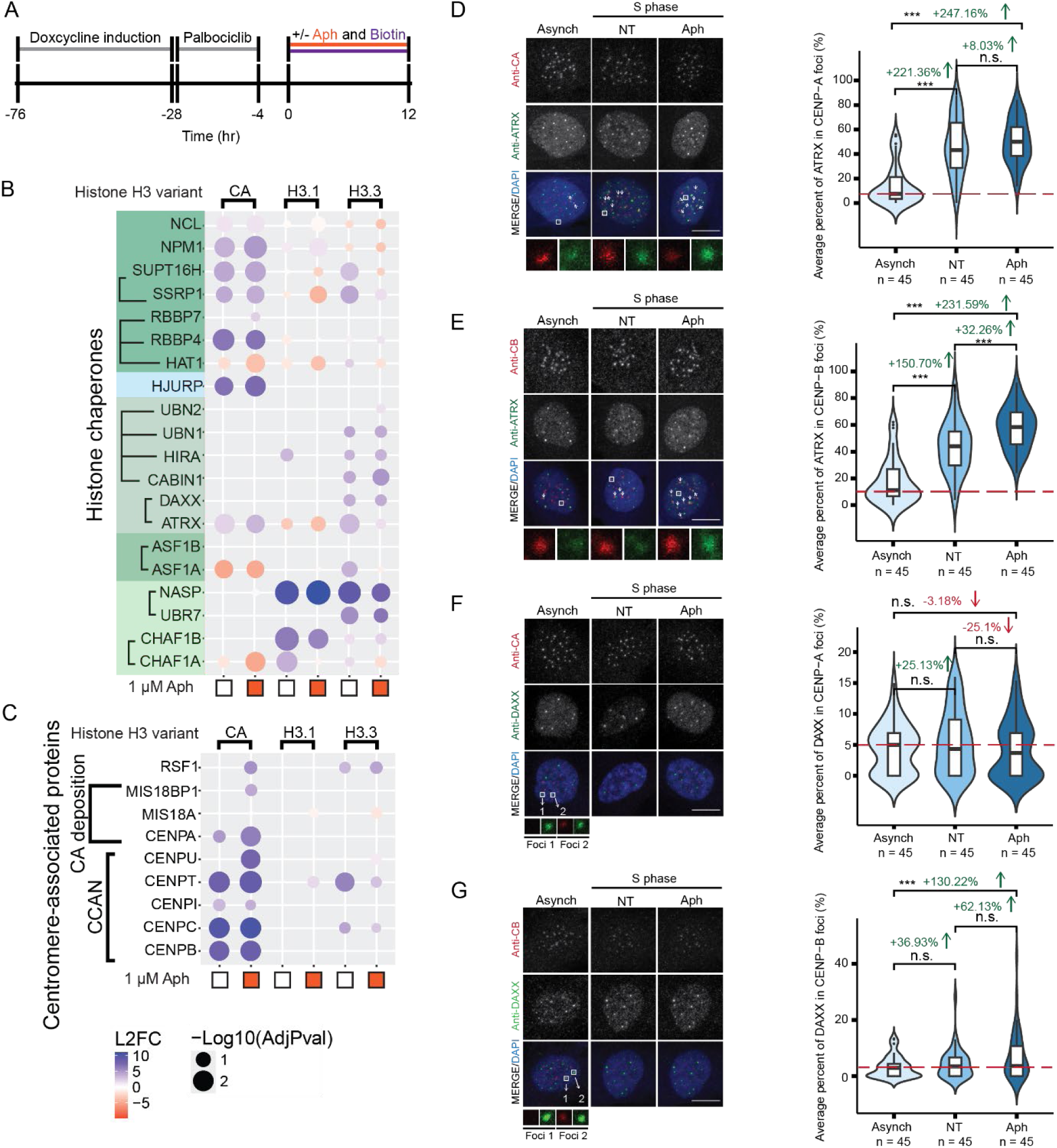
Replication stress leads to accumulation of ATRX to centromeres. **A)** Cell synchronization and replication stress workflow of Chromatin-TurboID experiments. **B)** Bubble plots representing pulldowns of histone-dependent interactions by proximity labeling under normal replication and replication stress (*N* = three biological replicates per condition). Protein groups were classified based on GO-term defined pathways including **B)** Histone chaperones and **C)** Centromere-associated. Colors represent Log_2_ Fold Change (L2FC) is of [histone/LaminA control]. -Log_10_ values are based on adjusted *p* values that are calculated by bonferroni correction test. **D)** Representative images of ATRX accumulation (left) and quantification of co-localization (right) at CENP-A or E) CENP-B. **F)** Representative images of DAXX accumulation (left) and quantification of co-localization (right) at CENP-A or at **G)** CENP-B. 15 cells per condition per biological replicate were measured. Not significant (n.s.) = *p* > 0.05, *p* ≤0.05, *p* ≤0.01 = **, *p* ≤0.001 = ***. Scale bar = 10 μm.

### ATRX is a centromere-associated protein and its association with CENP-A is replication-dependent manner

ATRX is best known for its interaction with H3.3 chaperone DAXX, where DAXX-ATRX complex deposits histone H3.3 at repetitive regions such as pericentromeric heterochromatin and telomeres (73,76,91). Our BP-MS findings revealed that CENP-A can associate with ATRX in a replication-dependent manner **(Figure 2E and 3B)**. ATRX is implicated to interact with CENP-A through DAXX chaperone under overexpression conditions, where DAXX-ATRX complex mediate ectopic CENP-A deposition at non-centromeric regions (53–55). Interestingly, our results did not show the presence of DAXX under replication conditions **(Figure 3B)**. However, several reports have implicated several mechanisms of centromere recruitment of ATRX and DAXX involving CENP-B and CENP-C have been reported (63,84,92,93). We thus asked whether ATRX and DAXX are recruited to centromeres during DNA replication and under stress. We hypothesize that replication stress-dependent loss of CENP-A is mediated by DAXX-ATRX.

To determine if ATRX is associated with centromeric chromatin, we examined the co-localization of ATRX to CENP-A and relative to centromere protein B (CENP-B) in both asynchronous and S-phase synchronized cells **(Figure 3D and E)**. We observed accumulation of ATRX at centromeres during DNA replication, but with significant elevation under replication stress condition **(Figure 3D and E)**. We observed both ATRX foci that are both positive in co-localized CENP-A or CENP-B and a subset of ATRX foci that are not co-localized with CENP-A or CENP-B foci **(Figure 3D)**. The presence of ATRX at both centromeric and non-centromeric regions highlights the diverse functions of ATRX. DAXX is a prominent interaction partner of ATRX; thus, we measured accumulation of DAXX to centromeres. Under the same conditions, we did not observe DAXX accumulation at centromeres through CENP-A or CENP-B **(Figure 3F and G)**. The lack of DAXX at centromeres is further supported by previous work which suggests DAXX is sequestered at PML nuclear bodies during interphase and is recruited to centromeres under ATR inhibition (27). However, siRNA-depletion of DAXX under the same conditions shows remarkable loss of ATRX at both general chromatin and centromeres, thus suggesting that DAXX is necessary for ATRX recruitment to centromeres **(SI Figure 5)**. Together these finds suggest that ATRX accumulate to centromeres in a DAXX-dependent manner in response to replication stress.

### ATRX and DAXX are necessary for stable inheritance of CENP-A

Centromere replication occurs in mid to late S-phase and co-occurs with the replication of heterochromatin (26,94–97). We asked whether ATRX or DAXX depletion influenced CENP-A and H3.3 levels by conducting siRNA depletion of ATRX and DAXX under asynchronous cells **(SI Figure 6A and B)**. Neither ATRX or DAXX depletion have effects on the expression of CENP-A or H3.3 **(SI Figure 6C-F)**. We next asked whether ATRX and DAXX are necessary for CENP-A retention during DNA replication. To address this question, we conducted siRNA depletion of either ATRX or DAXX under normal DNA replication in late-replicating RPE1 cells. To define late-replicating cells, nocodazole-arrested cells were pulse labeled with EdU **(Figure 4A)**. Depletion of either ATRX or DAXX in DNA replication was associated with loss of CENP-A **(Figure 4A and B)**. Conversely, the depletion of ATRX or DAXX in combination of Aph treatment did not result in additive effects of CENP-A loss **(Figure 4A and B)**. This finding suggests that ATRX or DAXX-mediated CENP-A loss may be also due to replication stress. Interestingly, DAXX removal in combination with Aph treatment rescues the loss of CENP-A, suggesting that DAXX function may oppose replication stress and contribute to CENP-A depletion when replication stress occurs via other mechanisms.

**Figure 4:**
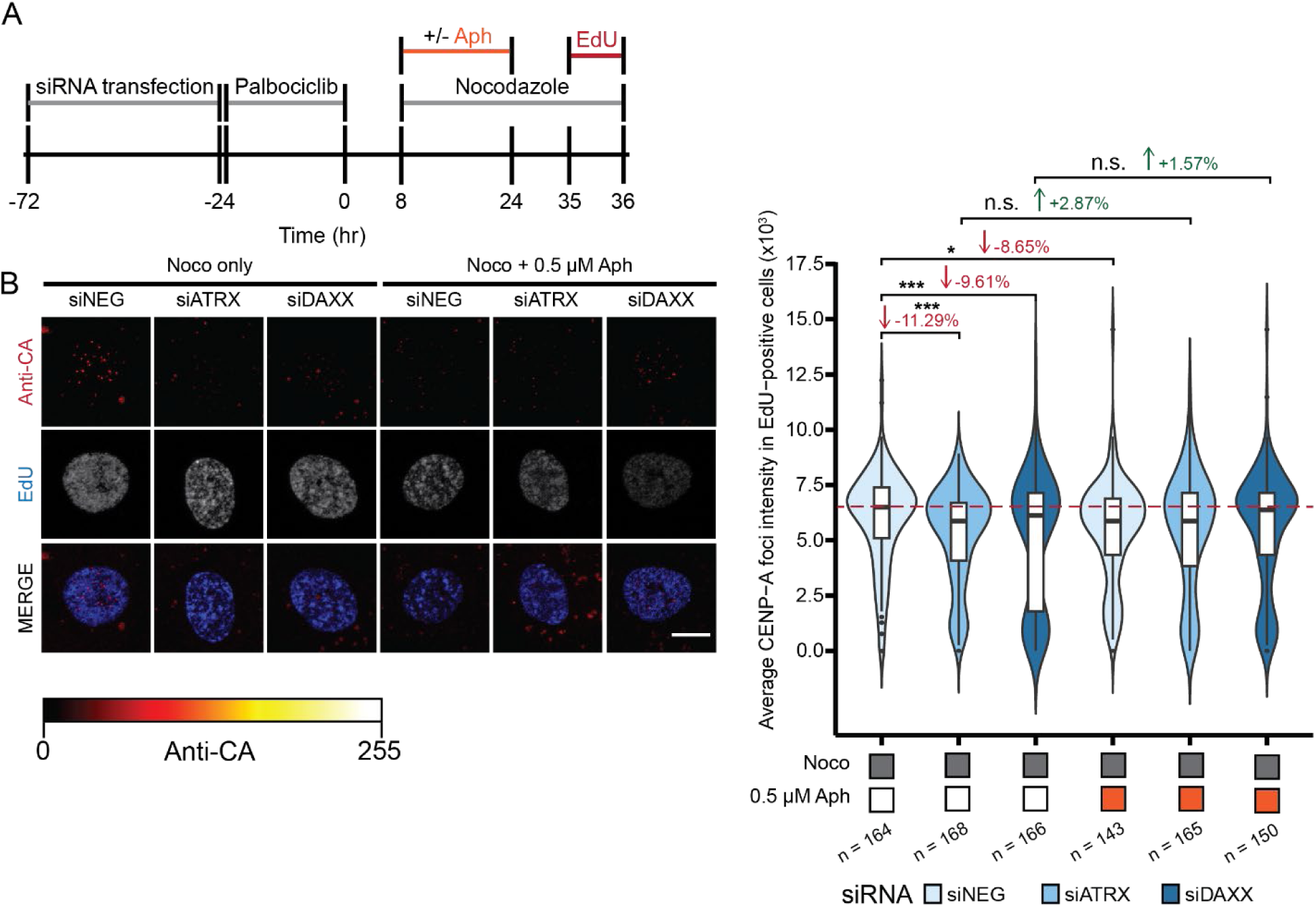
DAXX-ATRX depletion leads to specific loss of centromeric CENP-A. **A)** Graphical representation of experimental workflow. **B)** Representative images of late-replicating RPE1s with siRNA knockdown of ATRX or DAXX in RPE1 cells under control or Aph-induced stressed conditions (left) and quantification of CENP-A levels in EdU-positive cells (right). CENP-A alone is presented as an intensity gradient. Merged images represent CENP-A as red. Experiments were performed with three independent biological replicates and *n* = represents number of EdU-positive cells (approximately 50 cells measured per condition). Not significant (n.s.) = *p* > 0.05, *p* ≤ 0.05 = *, *p* ≤0.01 = **, *p* ≤0.001 = ***. Scale bar is 10 μm.

Our BP-MS results show that under Aph-induced stress, proximity labeling by CENP-A-TurboID resulted in the enrichment of DNA damage response protein 53BP1 **(SI Figure 4)**. We next asked whether the loss of CENP-A under these conditions would stimulate DNA damage repair at late S-phase **(Figure 5A)**. To address this, we measured both the global and centromeric levels of DNA damage response markers γH2AX and 53BP1. As expected, the depletion of either ATRX and DAXX depletion resulted in increased global levels of γH2AX **(Figure 5B)** and may reflect their additional roles in DNA damage response including R loop formation, homologous recombination, and replication fork restart (79–81). In addition, we observed elevated levels of centromeric γH2AX in either ATRX or DAXX-depleted conditions, which suggests a centromere-specific function and is independent of Aph-treated conditions. Similarly, we observed elevated global levels of the 53BP1 and at centromeres in response to Aph-induced stress relative to control **(Figure 4C)**. This observation in agreement with previous work that demonstrated recruitment of 53BP1 in response to rapid degradation of CENP-A (23). However, the depletion of ATRX or DAXX alone under normal replication did not increase global or centromeric levels of 53BP1 **(Figure 4C)**. This result contrasts with previous work that demonstrate enhanced 53BP1 recruitment to spontaneous DNA damage at centromeres as a response to shRNA depletion of DAXX (28).

**Figure 5:**
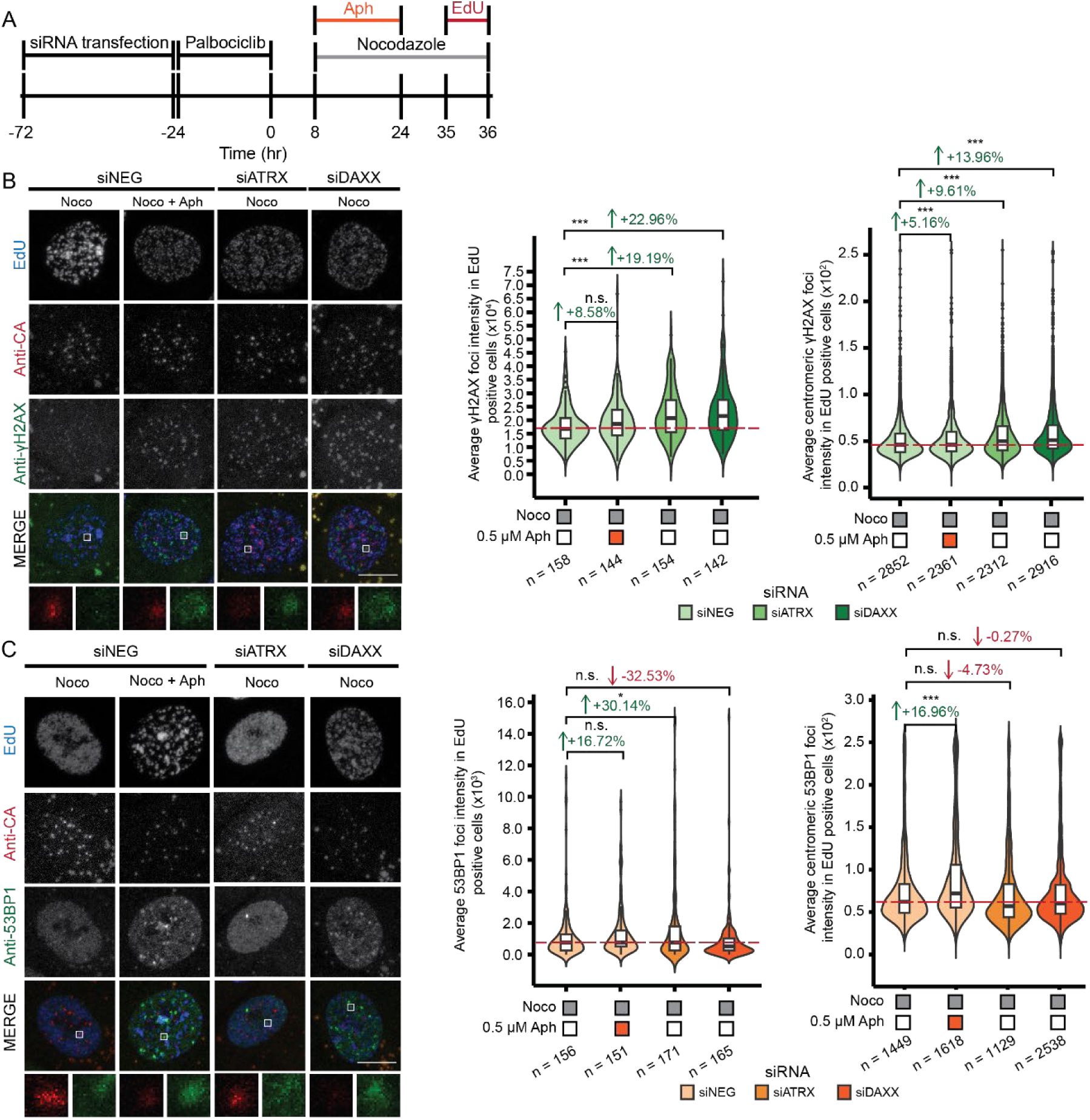
Replication stress leads to accumulation of DNA damage response factors 53BP1 and γH2AX at centromeres. **A)** Experimental scheme. **B)** Representative images of siRNA depletion of negative control (siNEG), ATRX, or DAXX in late S-phase RPE1 cells co-stained with CENP-A and γH2AX (left). Quantification of total γH2AX foci intensity (middle) and at centromeres using CENP-A (right). **C)** Representative images of late S-phase RPE1 under siRNA depletion conditions in Panel A, and co-stained with CENP-A and 53BP1 (left). Quantification of total 53BP1 foci intensity (middle) and at centromeres using CENP-A (right). For quantification of either γH2AX or 53BP1 approximately 50 cells were measured per condition. For quantification of nuclear γH2AX or 53BP1, *n* = number of cells measured. For nuclear 53BP1, outliers above 10000 were not shown but remained in data analysis. For centromeric quantification of either γH2AX or 53BP1, *n* =number of centromeres measured. Dashed line indicates baseline average intensity relative to noco only siNEG control. Not significant (n.s.) = *p* > 0.05, *p* ≤0.05, *p* ≤0.01 = **, *p* ≤0.001 = ***. Experiments were performed three independent biological replicates. Scale bar is 10 μm.

Replication stress has been shown to induce global heterochromatin formation (98,99). We next asked whether replication stress increases heterochromatin at centromeres. To address this, we measured the colocalization of heterochromatin modification H3K9me3 at centromeres under Aph-treated conditions. Under Aph-treated conditions we observed a two-fold increase of centromeric H3K9me3 in late S-phase cells relative to no treatment control **(SI Figure 7)**. This result suggests that heterochromatin formation at centromere occurs in response to replication stress.

We next asked whether ATRX, or DAXX-dependent loss of CENP-A persists into the next G1 of subsequent cell cycle. We hypothesize that CENP-A loss during DNA replication results in lower levels of CENP-A in subsequent G1. To address this, we synchronized ATRX or DAXX-depleted RPE1 cells at G1 by palbociclib. We subsequently released the cells into S phase and arrested them in the next G1 phase **(Figure 6A)**. Neither ATRX nor DAXX depletion interfere with proper cell cycle progression into the next G1 **(SI Figure 8)**. We observed lower levels of CENP-A under conditions of ATRX and DAXX depletion relative to control in G1-arrested cells **(Figure 6C)**. Similarly, we observed lower levels of CENP-A under conditions of Aph-induced stress. Unlike in late-S phase, under ATRX-depleted conditions in combination with Aph treatment, we see an additive effect on CENP-A loss in the subsequent G1 **(Figure 6C)**. Like replication stress in late S-phase, we observed a consistent phenotype in which DAXX removal in combination with Aph treatment rescues the loss of CENP-A in G1 **(Figure 6C)**.

**Figure 6:**
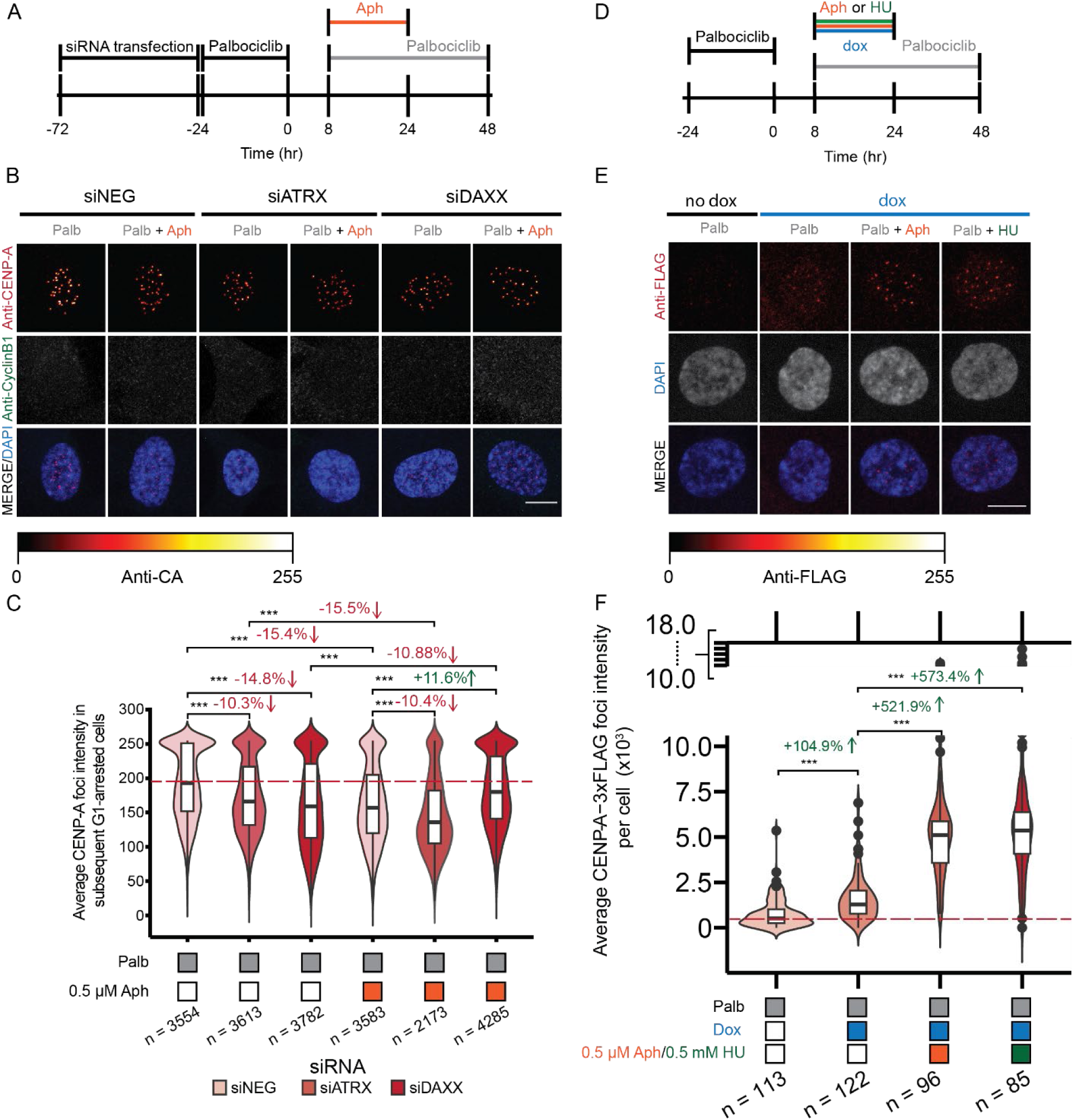
Replication stress leads to loss of parental CENP-A in subsequent cell cycle phase. **A)** Schematic of experimental design. **B)** Representative images of siRNA depletion of negative control (siNEG), ATRX, or DAXX in late S-phase RPE1 cells. **C)** Quantification of total CENP-A levels at post-replicative G1-arrested cells. **D)** Schematic of experimental design of FLAG-tag monitoring of parental CENP-A. **E)** Representative images of CENP-A-3xFLAG under replicating or stressed conditions in RPE1 cells. **F)** Quantification of CENP-A-3xFLAG levels (right). CENP-A or FLAG alone is presented as an intensity gradient. Merged images represent CENP-A or FLAG as red. Dashed line indicates baseline average intensity relative to no aphidicolin (Aph) control. n = represents number of cells measured per group. Not significant (n.s.) = *p* > 0.05, *p* ≤0.05 = *, *p* ≤0.01 = **, *p* ≤0.001 = ***. Experiments were performed with three independent biological replicates, except in **F)** which was performed with two independent biological replicates. Total percentage of cells in G1, S, or G2/M phase under siRNA depleted conditions. Error bars represent standard error of mean.

To determine whether loss of CENP-A was a result of *de novo* deposition or retention during replication, we generated a stable cell line that expressed FLAG-tagged CENP-A under a doxycycline inducible promoter, which allows for the monitoring of CENP-A in a pulse chase manner **(SI Figure 9)**. With our inducible FLAG-tagged CENP-A system, we measured FLAG-tagged CENP-A levels under conditions of replication stress induced by either Aph or HU treatment in cells synchronized in the subsequent G1 phase **(Figure 6D)**. Under these conditions, we observed increased levels of CENP-A in cells treated with either Aph or HU relative to control **(Figure 6E)**. In addition, we observed fewer cells with deposited FLAG-tagged CENP-A in the absence of replication stress. This suggests that CENP-A retention is negatively influenced by replication stress. As such we rationalize that increases in deposited FLAG-tagged CENP-A under replication stress conditions are due to the loss of endogenous CENP-A retention allowing for new deposition to occur in G1. Altogether, these findings suggests that CENP-A retention is susceptible to replication stress and can lead to loss of centromere identity.

## 4. Discussion

Parental CENP-A is stably retained by the HJURP chaperone and MCM2 helicase during centromere replication in late S-phase where it is redistributed onto newly synthesized centromere chromatin along with CCAN components (14,94). In this study we show that the fidelity of this process of CENP-A nucleosome re-assembly during DNA replication is sensitive to replication stress induced by pharmacological inhibitors targeting independent pathways in DNA metabolism. The overall effect of replication stress leads to reduced retention of parental CENP-A during S-phase that persists into G2 **(Figure 2)**. This suggests that cells are unable to recover CENP-A nucleosome loss during these times in the cell cycle, consistent with the restriction of new CENP-A nucleosome distribution to G1-phase (100,101). Inhibition of DNA replication by directly affecting DNA polymerase function (aphidicolin) or limiting nucleotide pools (hydroxyurea) inhibits the addition of new nucleotides but does not alter MCM helicase function (102,103). This results in the uncoupling of the MCM helicase complex from the replication fork and increased amounts of intervening single-stranded DNA (102,103). Since MCM2 contributes to the retention of both canonical and centromeric nucleosome retention, this suggest that coupling of the MCM helicase complex to the replication fork is critical for efficient recycling of histones (14–18).

Under normal replication conditions, the unoccupied regions left by redistribution of the existing centromeric nucleosomes into sister centromeres are filled by histone H3.3 in S-phase (19). These are subsequently removed for the new assembly of CENP-A in G1 phase (19). We observe increased levels of methylated H3K9 at centromeres following replication stress, suggesting that more H3 may occupy the centromere when CENP-A redistribution in S-phase fails. Interestingly, depletion of DAXX rescued the effect of CENP-A loss from aphidicolin treatment suggesting that CENP-A may compete with H3.3 in redeposition at the centromere. Our work suggests the importance of H3.3 deposition mediated by DAXX-ATRX to serve as placeholders during unperturbed replication of centromeres for the loading of new CENP-A in the subsequent G1 phase **(Figure 7)** (3,19,104).

**Figure 7:**
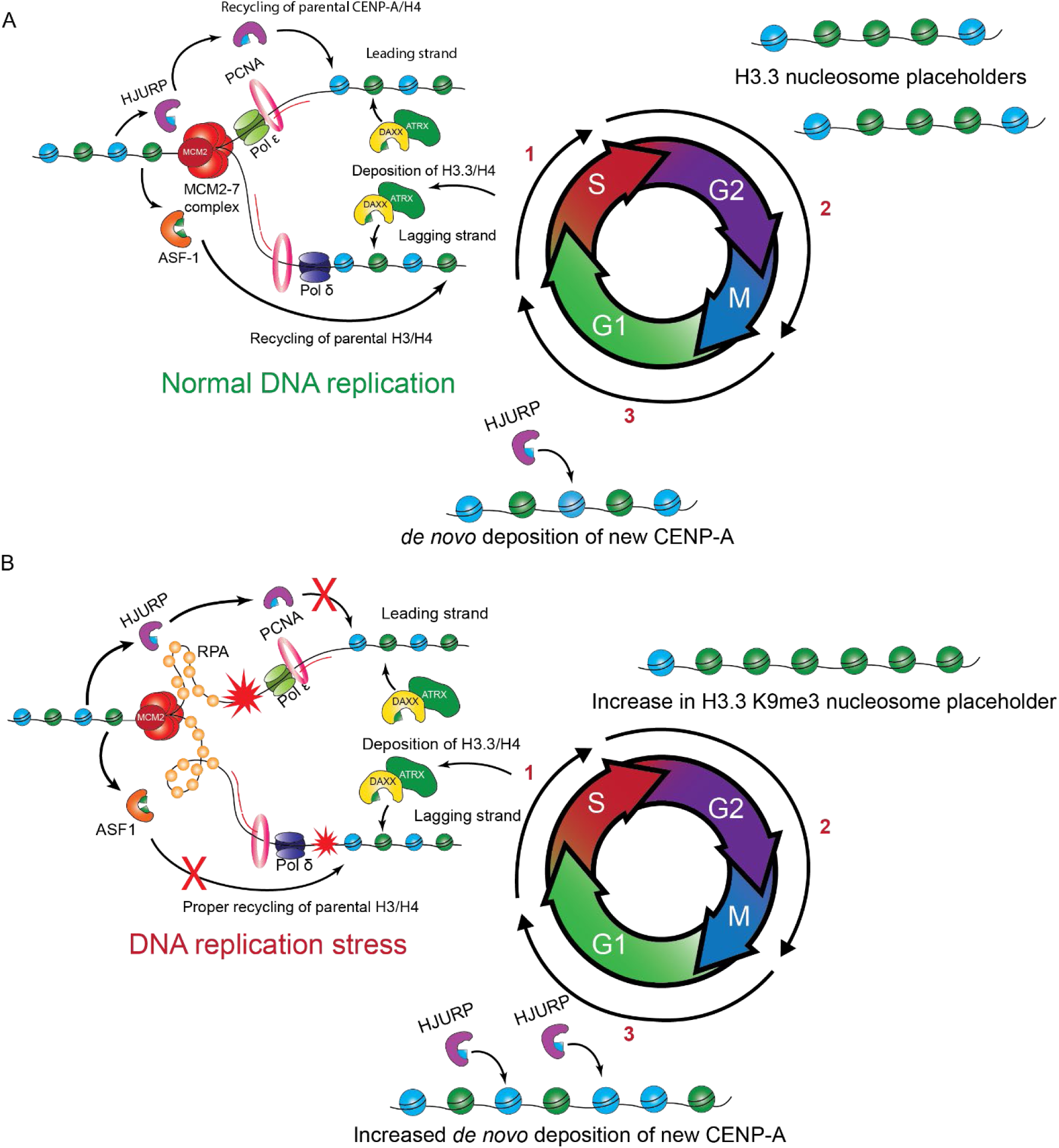
Centromere identity is susceptible to replication stress independent of underlying mechanism. **A)** During normal centromere replication, CENP-A nucleosomes are retained and redistributed onto newly synthesized DNA mediated by HJURP chaperone and MCM2-7 complex. H3.3 nucleosomes are deposited by DAXX-ATRX into newly replicated centromeres serving as placeholders for loading of new CENP-A nucleosomes in subsequent G1. **B)** DNA replication stress leads to heterochromatin formation in a DAXX-ATRX dependent manner and reduced parental CENP-A retention. In the following G1 phase, a compensatory effect occurs where new deposition of CENP-A is increased to overcome loss of parental CENP-A resulting from replication stress.

In addition to the pharmacological effects of replication inhibitors, we show that depletion of ATRX or DAXX also limits the retention of CENP-A nucleosomes. This could be a result of ATRX and DAXX function in preventing genomic instability. In the context of replication stress at heterochromatin regions, ATRX and DAXX are recruited to stalled replication forks where they mediate H3.3-specific deposition and recruit downstream DNA response factors to stabilize replication forks (77,80,81,105). Previous work by single DNA molecule fiber analysis has demonstrated the accumulation of H3K9me3 deposition at stalled replication forks within 1 hour when induced by HU (98). Our proteomics analysis shows a unique proximal interaction and enrichment of CENP-A and ATRX under conditions of normal replication and Aph-induced stress. This suggests that under normal conditions, ATRX responds to replication stress occurring at the centromere, which is enhanced upon replication stress. The enhanced H3.3 deposition association with recruitment may in turn limit CENP-A redeposition.

Despite the loss of existing CENP-A during DNA replication, when we measured new deposition in the next G1, cells treated with either aphidicolin or hydroxyurea showed an increased recruitment of new CENP-A. These data suggest that cells compensate for the CENP-A during a challenging round of DNA replication by depositing additional CENP-A in the following G1. CENP-A occupies only a small fraction of the alpha-satellite repeats within the centromere. H3K9me2 has been proposed to limit the distribution of CENP-A within the centromere (65,106), but these regions may also provide a yardstick to define how much CENP-A is deposited per centromere.

The repetitive nature of centromere regions is a constant challenge for the replication of centromeric DNA and the epigenetic inheritance of centromeres across cell generations (20,23,26,107). The repetitive regions are sources for DNA recombination events, and the formation of DNA secondary structures that perturb proper progression of replication machinery (1,20,21,28). Previous studies have demonstrated CENP-A and its associated CCAN components CENP-C and CENP-T to protect these susceptible regions (23,40,107). Human centromeres depleted of these components are more prone to exacerbated types of chromosome segregation defects include micronuclei formation, chromosome breakages, and anaphase bridges (23,40,107). Therefore, the retention of CENP-A nucleosomes may be critical not only for accurate segregation of chromosomes, but also for suppressing recombination events within the centromere repeats that contribute to chromosome rearrangements.

Taken together, the findings of our work demonstrate the link between replication stress and its effects on the fidelity of CENP-A-containing nucleosomes during the replication of centromeres.

## 5. Materials and Methods

### Cell culture and transfections

hTERT-RPE1 cells were grown and maintained in DMEM/F12 (Gibco, 11320033) medium supplemented with 10% (v/v) FBS (Biotechne, S10350) and 1% (v/v) Penicillin and Streptomycin (Sigma, P0781-100ML) at 37°C with 5% CO_2_. siRNA transfections were performed using Lipofectamine RNAiMAX transfection reagent (Invitrogen, 13778150) as per manufacturer’s protocol. The following siRNAs were used: siNEG control (Ambion: 4390846), siATRX (Horizon Discovery: L-006524-00-0005), and siDAXX (Thermo Fisher Scientific: s30814).

### Stable cell lines

TurboID constructs were generated using pCW57.1 vector (Addgene: 41393) by Hi-Fi DNA assembly (New England Biolabs, E2621S) at insertion sites between NheI and AgeI. Coding sequences of *CENPA, H3C1, H3-3A, TURBOID, and LMNA* were PCR amplified from gene blocks (IDT). For the generation of CENP-A-3x-FLAG cell line, the *CENPA* coding sequence was PCR amplified and cloned into pCW57.1 vector containing C-terminal 3xFLAG tag sequence using HiFi DNA assembly (New England Biolabs, E2621S). In brief, stable cell lines were generated using lentiviral transduction using Lipofectamine 3000 transfection reagent (Invitrogen, L3000001) as per manufacturer protocol. After transduction, positive cells were puromycin selected. Monoclonal cells of CENP-A overexpressing cells were isolated by single cell dilution. All cell lines were screened by immunofluorescence staining for proper expression and cellular localization. Additionally, proper expression of monoclonal cell lines was validated by immunoblot.

### Cell synchronization and drug treatments

hTERT-RPE1 cells were plated at 30% confluency relative to surface area of tissue culture dishes. In general, cells were treated with 300 nM Palbociclib (Sigma, PZ0383-5MG) for 24 hours for synchronization at G1 phase. To release from G1-arrest, palbociclib-media was removed, and cells were washed three times with 1X DBPS (Gibco, 14-040-133) and fresh DMEM/F12 was replenished. Cells were treated either 0.5 µM or 1 µM Aphidicolin (Aph) (Cayman Chemical, 14007) or with 0.5 mM or 1 mM Hydroxyurea (HU) (Sigma, H8627-5G). For G2/M-arrested cells, cells were treated with 50 ng/mL of nocodazole. Details pertaining treatment times are specified in figure captions.

### DNA content analysis by flow cytometry

Cells were harvested using 3 mM EDTA in 1X PBS. Cells were centrifuged at 1200 RPM for 5 mins, and washed once with 1X DPBS (Gibco, 14190144). Cells were resuspended in 200 µL of 1 X DPBS, fixated with 5 mL of 70% ethanol, and stored at 4°C. Ethanol-fixed cells were centrifuged at 1600 RPM for 5 mins, and washed with 1 mL of 1X DPBS + 5% FBS (Biotechne, S12450). Cell pellets were resuspended in 1 mL of propidium iodide/RNAseA solution [10 µg/mL propidium iodide (Sigma, P4170-10MG) 250 µg/mL RNase A in PBS (Sigma, 10109169001b) + 1% (v/v) FBS)] and incubated at 37°C for 1 hour prior to flow cytometry. FACs analysis and cell cycle profiles were performed on FloJo (v10.10.0).

### Immunoblots

Cell pellets were resuspended in RIPA (25 mM Tris-HCl, pH 7.6, 150 mM NaCl, 1% NP-40, 1% sodium deoxycholate, 0.1% SDS) supplemented with 1X protease inhibitor cocktail (Sigma, 11873580001), 2 mM MgCl_2_ and 1 µL/mL of lysis buffer of Universal Nuclease (25 kU) (Thermo Fisher Scientific, 88701), and incubated at room temperature for 15 mins. Cell lysates were centrifuged at 13,000 x *g* for 10 mins at 4°C, and the resulting clarified supernatants were incubated with pre-equilibrated Dynabeads MyOne Streptavidin C1 (Thermo Fisher Scientific, 65001) on end-over-end rotator for overnight at 4°C. Beads were pelleted by centrifugation at 1000 RPM for 1 min. Beads were washed twice with 1 mL RIPA lysis buffer followed by once with 1 mL RIPA supplemented with 2% SDS. Beads were subsequently washed once with 1 mL of 1 M KCl followed by 1 mL of 0.1 M NaCO_3_. Beads were washed with 1 mL of 2 M Urea in Tris-HCl pH 8.0 followed by two washes of 1 mL of RIPA lysis buffer. Beads were boiled in 60 µL of 1x Laemmli SDS sample buffer + 2 mM Biotin at 95°C for 10 mins. Resulting elutants were used for immunoblot detection.

Immunoblotting of lysates and pulldowns were conducted using the following primary antibodies: Anti-CENP-A (Cell Signaling Technology, 2186S, 1:1000), Anti-ASF1A (Cell Signaling Technology, 2990S, 1:1000), Anti-ATRX (Invitrogen, PA551656, 1:1000), Anti-DAXX (Invitrogen, MA1-19731, 1:1000), Anti-FLAG (Sigma Aldrich, F1804-50UG, 1:1000), Anti-GAPDH (Invitrogen, PA1-16777, 1:5000), Anti-Histone H3.3 (Invitrogen, MA5-24667, 1:1000), Anti-HA-HRP tag (Cell signaling technologies, 2996S, 1:1000), Anti-HJURP (Custom antibody, 1:1000), Streptavidin-HRP (Jackson ImmunoResearch, NC9705430, 1:10000), Anti-Ras (Cell Signaling Technology: 91054S). Primary antibodies were incubated in 5% (w/v) milk in TBST blocking buffer for overnight at 4°C. Blots were washed with TBST wash buffer four times at 15 mins each. Secondary antibodies include Anti-Mouse-HRP (Jackson ImmunoResearch, 111-035-003) and Anti-Rabbit-HRP (Jackson ImmunoResearch, 111-035-003). Blots were incubated in secondary antibodies for 1 hour at room temperature, and subsequently washed 4 times at 15 mins each prior to chemiluminescence imaging.

### Immunofluorescence staining

Cells were straight fixed onto coverslips with 4% (v/v) paraformaldehyde in 1X DPBS solution for 10 mins, and subsequently quenched in 100 mM Tris (pH 7.5) for 5 mins. Fixed cells were pre-extracted with 0.1% (v/v) triton-X-100 in PBS for 3 mins and incubated in blocking solution [0.1% (v/v) triton X-100 in PBS, 0.2% (v/v) BSA, 2% (v/v) FBS] for 1 hour. Cells were incubated with indicated primary antibodies for 1 hour. Primary antibodies include the following: Anti-ATRX (Invitrogen, PA551656, 1:500), Anti-CENP-A (Invitrogen, MA1-20832, 1:250), Anti-CENP-A (Cell Signaling Technology, 2186S, 1:1000), Anti-CENP-T (Custom antibody, 1:5000), Anti-Cyclin B1 (Cell Signaling Technology: 12231S, 1:1000), Anti-DAXX (Invitrogen, MA1-19731, 1:1000), Anti-FLAG (Sigma-Aldrich, F180450UG, 1:1000), Anti-HA tag (Cell Signaling Technology, 2367S, 1:1000), Anti-γH2A.X (Cell Signaling Technology, 2577S, 1:1000), Anti-H3K9me3 (AbCam: ab8898-100ug), Anti-p21 (Cell signaling technology: 2947, 1:500). Cells were washed with 1X PBS three times, and subsequently incubated with indicated secondary antibodies. Secondary antibodies include the following: Cy3-Streptavidin (Jackson ImmunoResearch, 016-160-084, 1:10000), Goat Cy3-Anti-Mouse (Jackson ImmunoResearch, 115-165-146, 1:1000), and Donkey Anti-AlexaFluor488 (Jackson ImmunoResearch, 711-545-152, 1:1000). DNA was stained using DAPI solution (1 µg/mL of DAPI in 1X PBS), and coverslips were mounted onto cover slides using ProLong Gold Antifade (Thermo Fisher Scientific, P36930). Coverslips were left to dry overnight at room temperature prior to imaging.

### EdU treatment and labeling

The Click-iT cell proliferation assay (Invitrogen, C10340) using AlexaFluor 647 to label newly synthesized DNA. In brief, late-S phase cells were treated with 10 µM EdU one hour prior to straight-fixation for immunofluorescence staining as described above. Click-chemistry reaction using AlexaFluor 647 was performed as per manufacturer protocol.

### Imaging and Image Analysis

All image acquisition of fixed cells was performed using an oil-immersion 64X objective lens on LSM800 confocal microscope (Zeiss). Displayed representative images are maximum stacked images. For image analysis of centromeric CENP-A or CENP-A-FLAG foci, intensities were measured from raw stack images using Centromere recognition and Quantification (CrAQ) plugin with CENP-A or CENP-T as a centromere reference marker (108). Similarly, for measurement of centromeric γH2AX and 53BP1 foci, intensities were from raw stack images using CENP-A as a centromere reference marker. For image analysis of nuclear CENP-A, γH2AX, and 53BP1 foci in S-phase positive cells, intensities were measured from stacked images of EdU-positive nuclei masks from a custom script in ImageJ/Fiji. Quantification of centromeric H3K9me3 was performed using CrAQ using CENP-A as centromere reference marker in EdU-positive cells. For image analysis of nuclear CENP-A foci in G1-phase cells, intensities were measured from maxima using a prominence tolerance value of 80 from stacked images of nuclei masks from a custom script in ImageJ/Fiji. Statistical analysis was applied using student’s *t* test. Image channels were scaled identically within panels of figures.

### Gene expression analysis by quantitative reverse-transcriptase PCR

Total RNA was extracted using RNEasy RNA extraction kit (Qiagen: 74106) as per manufacturer protocol. cDNA was prepared from 2 µg of total extracted RNA using the iScript cDNA Synthesis kit (Bio-Rad, 17088900). RT-qPCR was performed using 10-fold diluted cDNA, and iTaq universal SYBR green supermix (Bio-Rad, 1725121), and forward and reverse primers to a final concentration of 500 nM. RT-qPCR was performed as described in iTaq SYBR manual. Gene expression analysis was performed using 2-ΔΔCt method. Three technical replicates were analyzed per every biological replicate, and experiments were performed with N = three independent biological replicates. The study used following qPCR primers used: *ATRX* (Forward: ACGGCGTTAGTGGTTTGTCCTC, Reverse: GCAGCATGTAGCTTCTCTCCTG), *CENPA* (Forward: CTCGTGGTGTGGACTTCAAT, Reverse: CTGCATGTAAGG TGAGGAGATAG), *DAXX* (Forward: GAAGCCTCCTTGGATTCTGGTG, Reverse: CATCACTCTCCTCATCGTCTTCG), *GAPDH* (Forward: GTCTCCTCTGACTTCAACAGCG, Reverse: ACCACCCTGTTGCTGTAGCCAA), *H3F3A* (Forward: GGGTGAAGAAACCTCATCGT, Reverse: GGAAGTTTGCGAATC).

### Proximity labeling by Chromatin-TurboID

Four 15 cm dishes of cells were prepared per condition at confluency of 50% to prevent contact inhibition. Cell lines were treated with 0.5 µg/mL of doxycycline (Sigma, 50-165-6938) for 48 hours. Cells were synchronized to G1 for 24 hours with 300 nM Palbociclib (Sigma, PZ0383-5MG). Cells were washed three times with 1X DBPS (Gibco, 14-040-133) and fresh DMEM/F12 was replenished. Cells were treated with 1 µM Aphidicolin (Aph) (Cayman Chemical, 14007) and 100 µM biotin (Sigma, B4501-1G) for 8 hours prior to harvest. At designed time points as indicated in figure legends, cells were washed three times with 1X DBPS. Cell pellets were washed once with 1X DPBS and snap frozen.

Cell pellets were resuspended in RIPA (25 mM Tris-HCl, pH 7.6, 150 mM NaCl, 1% NP-40, 1% sodium deoxycholate, 0.1% SDS) supplemented with 1X EDTA-free protease inhibitor cocktail (Sigma, 11873580001), 2 mM MgCl_2_ and 1 µL/mL of lysis buffer of Universal Nuclease (25 kU) (Thermo Fisher Scientific, 88701), and incubated at room temperature for 15 mins. Cell lysate was centrifuged at 13,000 x *g* for 10 mins at 4C, and resulting clarified supernatants were incubated with pre-equilibrated Dynabeads MyOne Streptavidin C1 (Thermo Fisher Scientific, 65001) on end-over-end rotator for overnight at 4C. Beads were pelleted by centrifugation at 1000 RPM for 1 min. Beads were washed twice with 1 mL RIPA lysis buffer followed by once with 1 mL RIPA supplemented with 2% SDS. Beads were subsequently washed once with 1 mL of 1 M KCl followed by 1 mL of 0.1 M NaCO_3_. Beads were washed with 1 mL of 2 M Urea in Tris-HCl pH 8.0 followed by two final washes with 1 mL of RIPA lysis buffer. Beads were boiled in 60 µL of 1x Laemmli SDS sample buffer + 2 mM Biotin at 95°C for 10 mins.

### Sample preparation for biotin purification mass spectrometry

Samples were prepared by in-gel digestion using a 4-12% Bis-Tris SDS-PAGE (Invitrogen) stacking gel. Excised gel bands were destained with 50% ACN:50% 100 mM ammonium bicarbonate (AmBic). Samples were reduced using 10 mM DTT in 100 mM AmBic, and subsequently alkylated with 100 mM Iodoacetamide in 100 mM AmBic for 30 mins. Gel bands were treated with 200 ng of trypsin dissolved in 100 mM AmBic with gentle shaking at 37°C overnight. Supernatant was desalted using C18 spin columns (Thermo Fisher Scientific, 89851).

### Mass spectrometry analysis

LC-MS analysis of proximity labeled samples under asynchronous conditions was performed on Dionex UltiMate 3000 nanoLC (Thermo Fisher Scientific) coupled to Q-Exactive HF Orbitrap mass spectrometer (Thermo Fisher Scientific). Analytical separation was conducted using a 75 µm x 10.5 cm PicoChip column and packed with 3 μm ReproSil-Pur® C18 beads (New Objective). For each sample, 1-5 μL of sample was injected with a flow rate of 300 nL/min. Elution of samples performed on Dionex UltiMate 3000 nanoLC (Thermo Fisher Scientific) used a total 120 min gradient runtime between buffer A (0.1% formic acid and 99.9% Optima LC/MS grade water) and buffer B (80% Acetonitrile, 19.9% Optima LC/MS grade water, and 0.1% formic acid). The gradient used is as follows: 0% B at the beginning, 5% B at 5 min, 40% B at 100 min, 60% B at 104 mins, 90% B at 107-113 min followed by re-equilibration of 5% B at 113-120 min. MS parameters using Q-Exactive HF Orbitrap mass spectrometer was performed using an Ion Max source with HESI probe (Thermo Fisher Scientific). For electrospray ionization, positive static spray voltage was set at 2550 V. For full MS^1^, scan range was set to 375-2000 *m/z* with RF lens: 70%, Orbitrap resolution: 60,000, AGC Target: 3e^6^, Maximum injection time (ms): 120, Microscans: 1.

MS/MS data-dependent acquisition (DDA) by TopN was performed through fragmentation isolated precursor ions with charges between +2 to +5. The following parameters were used for DDA: scan range: 375-2000 *m/z*, Dynamic exclusion mode: exclusion duration (s): 20, mass tolerance: 5 ppm (low) and 5 ppm (high). Data dependent mode: cycle time (s): 2, Isolation window (*m/z*): 1.6, Normalized collisional energy = 30%, Orbitrap resolution 30,000, Define first mass (*m/z*): 110, AGC target: 1e^5^, Minimum AGC target: 1e^3^, Maximum injection time (ms): 120, Microscans: 1, Intensity threshold: 8.3e^3^.

LC-MS analysis of DNA replication and stressed samples was performed on Vanquish Neo UHPLC (Thermo Fisher Scientific) coupled to Orbitrap Exploris 240 mass spectrometer (Thermo Fisher Scientific). Analytical separation was conducted using a UHPLC C18 column (Ion Opticks, AUR3-15075C18-CSI). For each run, 1-5 μL of sample was injected. The flow rate was set at 0.2 µL/min. Elution of peptides from analytical separation column was performed using a 120 min gradient between buffer A (0.1% formic acid and 99.9% Optima LC/MS grade water) and buffer B (80% acetonitrile, 19.9% Optima LC/MS grade water, and 0.1% formic acid): 0% B at the beginning, 8% B at 1 min, 28% B at 86 min, 50% B at 106 min, 100% B from 107 to 110 min, 0% B from 111 to 120 Electrospray ionization using Orbitrap Exploris 240 was performed using an Nanospray Flex Ion Source (Thermo Fisher Scientific) and positive static spray voltage was set at 1800 V. For full MS^1^, scan range was set to 350-1600 *m/z* with Application mode: Peptide mode, RF lens: 60%, Orbitrap resolution: 120,000, Normalized AGC Target (%): 300, Maximum injection time (ms): 25, Microscans: 1, and Intensity threshold: 5.0e^3^.

Data-dependent acquisition (DDA) by TopN was performed through fragmentation isolated precursor ions with charges between +2 to +5. The following parameters were used for DDA: Dynamic exclusion mode: exclusion duration (s): 30, mass tolerance: 5 ppm (low) and 5 ppm (high). Data dependent mode: cycle time (s): 2, ddMS^2^ parameters isolation window (*m/z*): 1.5, Normalized collisional energy: 30%, Orbitrap resolution 15,000, scan range mode: define first mass, first mass (*m/z*): 200, Normalized AGC target (%): 100, Maximum injection time (ms): 50, Microscan: 1.

### Analysis of BP-MS data

The human FASTA database (UniProtProteome: UP000005640) used in this study was downloaded from UniProt. Database searching and processing of raw files were conducted with MaxQuant (v2.1.4.0) using default settings. MS data was normalized to total ion current, and adjusted *p* values were calculated based on Bonferroni correction. All mass spectrometry .raw files were deposited into MassIVE (MSV000097501).

### Data Visualization

Violin and volcano plots were visualized with Rstudio using libraries ggplot2 and EnhancedVolcano package (10.18129/B9.bioc.EnhancedVolcano), respectively. GO Enrichment analysis was performed by using Metascape under the Express Analysis function using Input Species: *Homo sapiens* (109). S-phase progression highlighted in **Figure 1B** was plotted using GraphPad (v10.2.2).

### Limitations of study

It is well established that CENP-A overexpression leads to its ectopic deposition into non-centromeric chromatin (53–55,110). In this study, we generated stably overexpressing CENP-A-TurboID cell line by lentiviral transduction. While CENP-A overexpression in this system is a concern, we screened for monoclonal cell populations, and validated centromeric localization of CENP-A-TurboID. In addition, our BP-MS results show a lack of H3.3 chaperones DAXX and HIRA, which are proposed to be the main drivers of ectopic CENP-A deposition under overexpression conditions. These findings give us confidence that any chromatin-associated proteins captured in CENP-A-TurboID cell line are not a result of overexpression.

## Supporting information

Lee_et_al_RS_CENPA_SUPP_INFO_initial_submission

## Acknowledgements

This work was supported by the National Institute of General Medical Sciences grants R01 GM 111907, R01 GM143638, and P41GM108569 at Northwestern University. A.S.L is a trainee fellow under the Chemistry of Life Processes Predoctoral Training Grant (5T32GM105538-10) at Northwestern University. Proteomics was partially performed by the Northwestern Proteomics Core Facility, generously supported by NCI CCSG P30 CA060553 awarded to the Robert H Lurie Comprehensive Cancer Center. We would like to thank Dr. Ewelina Zasadzinska and Kelvin Wong for their initial contributions to the development of this study. We would like to thank members from both the Foltz and Kelleher labs for their comments on the manuscript.

## Abbreviations

BP-MS: biotin purification mass spectrometry
CAP: chromatin-associated protein
CENP-A: centromere protein A
CENP-B: centromere protein B
CENP-C: centromere protein C
CENP-T: centromere protein T
HJURP: holliday junction recognition protein
ATRX: Alpha-Thalassemia Mental Retardation X-linked protein
DAXX: death domain-associated protein 6
Aph: aphidicolin
HU: hydroxyurea
Palb: palbociclib
Noco: nocodazole

## References

1. Saayman X, Graham E, Nathan WJ, Nussenzweig A, Esashi F. Centromeres as universal hotspots of DNA breakage, driving RAD51-mediated recombination during quiescence. Mol Cell [Internet]. 2023;1–16. Available from: 10.1016/j.molcel.2023.01.004

2. Foltz DR, Jansen LET, Black BE, Bailey AO, Yates JR, Cleveland DW. The human CENP-A centromeric nucleosome-associated complex. Nat Cell Biol. 2006;8(5):458–69.

3. Black BE, Jansen LET, Maddox PS, Foltz DR, Desai AB, Shah J V., et al. Centromere Identity Maintained by Nucleosomes Assembled with Histone H3 Containing the CENP-A Targeting Domain. Mol Cell. 2007;25(2):309–22.

4. Pesenti ME, Raisch T, Conti D, Walstein K, Hoffmann I, Vogt D, et al. Structure of the human inner kinetochore CCAN complex and its significance for human centromere organization. Mol Cell [Internet]. 2022 Jun;82(11):2113–2131.e8. Available from: https://linkinghub.elsevier.com/retrieve/pii/S1097276522003902

5. Yatskevich S, Muir KW, Bellini D, Zhang Z, Yang J, Tischer T, et al. Structure of the human inner kinetochore bound to a centromeric CENP-A nucleosome. Science (1979) [Internet]. 2022 May 20;376(6595):844–52. Available from: https://www.science.org/doi/10.1126/science.abn3810

6. Foltz DR, Jansen LET, Bailey AO, Yates JR, Bassett EA, Wood S, et al. Centromere-Specific Assembly of CENP-A Nucleosomes Is Mediated by HJURP. Cell [Internet]. 2009;137(3):472–84. Available from: 10.1016/j.cell.2009.02.039

7. Dunleavy EM, Roche D, Tagami H, Lacoste N, Ray-Gallet D, Nakamura Y, et al. HJURP Is a Cell-Cycle-Dependent Maintenance and Deposition Factor of CENP-A at Centromeres. Cell. 2009;137(3):485–97.

8. Nardi IK, Zasadzińska E, Stellfox ME, Knippler CM, Foltz DR. Licensing of Centromeric Chromatin Assembly through the Mis18α-Mis18β Heterotetramer. Mol Cell. 2016 Mar 3;61(5):774–87.

9. Fujita Y, Hayashi T, Kiyomitsu T, Toyoda Y, Kokubu A, Obuse C, et al. Priming of Centromere for CENP-A Recruitment by Human hMis18α, hMis18β, and M18BP1. Dev Cell. 2007 Jan;12(1):17–30.

10. Shuaib M, Ouararhni K, Dimitrov S, Hamiche A. HJURP binds CENP-A via a highly conserved N-terminal domain and mediates its deposition at centromeres. Proc Natl Acad Sci U S A. 2010 Jan 26;107(4):1349–54.

11. Jansen LET, Black BE, Foltz DR, Cleveland DW. Propagation of centromeric chromatin requires exit from mitosis. Journal of Cell Biology. 2007 Mar 12;176(6):795–805.

12. Massey DJ, Koren A. Telomere-to-telomere human DNA replication timing profiles. Sci Rep. 2022 Dec 1;12(1).

13. Erliandri I, Fu H, Nakano M, Kim JH, Miga KH, Liskovykh M, et al. Replication of alpha-satellite DNA arrays in endogenous human centromeric regions and in human artificial chromosome. Nucleic Acids Res. 2014 Oct 13;42(18):11502–16.

14. Zasadzińska E, Huang J, Bailey AO, Guo LY, Lee NS, Srivastava S, et al. Inheritance of CENP-A Nucleosomes during DNA Replication Requires HJURP. Dev Cell. 2018;47(3):348–362.e7.

15. Richet N, Liu D, Legrand P, Velours C, Corpet A, Gaubert A, et al. Structural insight into how the human helicase subunit MCM2 may act as a histone chaperone together with ASF1 at the replication fork. Nucleic Acids Res. 2015;43(3):1905–17.

16. Clément C, Almouzni G. MCM2 binding to histones H3-H4 and ASF1 supports a tetramer-to-dimer model for histone inheritance at the replication fork. Nat Struct Mol Biol. 2015;22(8):587–9.

17. Petryk N, Dalby M, Wenger A, Stromme CB, Strandsby A, Andersson R, et al. MCM2 promotes symmetric inheritance of modified histones during DNA replication. Science (1979). 2018;361(6409):1389–92.

18. Huang H, Strømme CB, Saredi G, Hödl M, Strandsby A, González-Aguilera C, et al. A unique binding mode enables MCM2 to chaperone histones H3-H4 at replication forks. Nat Struct Mol Biol. 2015;22(8):618–26.

19. Dunleavy EM, Almouzni G, Karpen GH. H3.3 is deposited at centromeres in S phase as a placeholder for newly assembled CENP-A in G phase. Nucleus. 2011;2(2):146–57.

20. Chardon F, Japaridze A, Witt H, Velikovsky L, Chakraborty C, Wilhelm T, et al. CENP-B-mediated DNA loops regulate activity and stability of human centromeres. Mol Cell [Internet]. 2022;82(9):1751–1767.e8. Available from: 10.1016/j.molcel.2022.02.032

21. Shih HT, Chen WY, Wang HY, Chao T, Huang H Da, Chou CH, et al. DNMT3b protects centromere integrity by restricting R-loop-mediated DNA damage. Cell Death Dis. 2022;13(6):1–14.

22. Mishra PK, Chakraborty A, Yeh E, Feng W, Bloom KS, Basrai MA. R-loops at centromeric chromatin contribute to defects in kinetochore integrity and chromosomal instability in budding yeast. Mol Biol Cell. 2021;32(1):74–89.

23. Giunta S, Hervé S, White RR, Wilhelm T, Dumont M, Scelfo A, et al. CENP-A chromatin prevents replication stress at centromeres to avoid structural aneuploidy. Proc Natl Acad Sci U S A. 2021;118(10):1–9.

24. Gaillard H, García-Muse T, Aguilera A. Replication stress and cancer. Vol. 15, Nature Reviews Cancer. Nature Publishing Group; 2015. p. 276–80.

25. Saxena S, Zou L. Review Hallmarks of DNA replication stress. Mol Cell [Internet]. 2022;82(12):2298–314. Available from: 10.1016/j.molcel.2022.05.004

26. Scelfo A, Angrisani A, Grillo M, Barnes BM, Muyas F, Sauer CM, et al. Specialized replication mechanisms maintain genome stability at human centromeres. Mol Cell. 2024;1–18.

27. Trier I, Black EM, Joo YK, Kabeche L. ATR protects centromere identity by promoting DAXX association with PML nuclear bodies. Cell Rep [Internet]. 2023;42(5):112495. Available from: http://www.ncbi.nlm.nih.gov/pubmed/37163376

28. Pinto LM, Pailas A, Bondarchenko M, Sharma AB, Neumann K, Rizzo AJ, et al. DAXX promotes centromeric stability independently of ATRX by preventing the accumulation of R-loop-induced DNA double-stranded breaks. Nucleic Acids Res [Internet]. 2023 Dec 1; Available from: http://www.ncbi.nlm.nih.gov/pubmed/38038252

29. Reverón-Gómez N, González-Aguilera C, Stewart-Morgan KR, Petryk N, Flury V, Graziano S, et al. Accurate Recycling of Parental Histones Reproduces the Histone Modification Landscape during DNA Replication. Mol Cell. 2018;72(2):239–249.e5.

30. Li N, Gao Y, Zhang Y, Yu D, Lin J, Feng J, et al. Parental histone transfer caught at the replication fork. Nature [Internet]. 2024 Mar 6; Available from: https://www.nature.com/articles/s41586-024-07152-2

31. Serra-Cardona A, Zhang Z. Replication-Coupled Nucleosome Assembly in the Passage of Epigenetic Information and Cell Identity. Trends Biochem Sci [Internet]. 2018;43(2):136–48. Available from: 10.1016/j.tibs.2017.12.003

32. Yu C, Gan H, Serra-Cardona A, Zhang L, Gan S, Sharma S, et al. A mechanism for preventing asymmetric histone segregation onto replicating DNA strands. Science (1979). 2018;361(6409):1386–9.

33. Groth A, Corpet A, Cook AJL, Roche D, Bartek J, Lukas J, et al. Regulation of replication fork progression through histone supply and demand. Science (1979). 2007;318(5858):1928–31.

34. Mejlvang J, Feng Y, Alabert C, Neelsen KJ, Jasencakova Z, Zhao X, et al. New histone supply regulates replication fork speed and PCNA unloading. Journal of Cell Biology. 2014;204(1):29–43.

35. Jasencakova Z, Scharf AND, Ask K, Corpet A, Imhof A, Almouzni G, et al. Replication Stress Interferes with Histone Recycling and Predeposition Marking of New Histones. Mol Cell. 2010;37(5):736–43.

36. Groth A, Ray-Gallet D, Quivy JP, Lukas J, Bartek J, Almouzni G. Human Asf1 regulates the flow of S phase histones during replicational stress. Mol Cell. 2005;17(2):301–11.

37. Li W, Yi J, Agbu P, Zhou Z, Kelley RL, Kallgren S, et al. Replication stress affects the fidelity of nucleosome-mediated epigenetic inheritance. PLoS Genet. 2017;13(7):1–28.

38. Dreyer J, Ricci G, van den Berg J, Bhardwaj V, Funk J, Armstrong C, et al. Acute multi-level response to defective de novo chromatin assembly in S-phase. Mol Cell [Internet]. 2024 Dec;84(24):4711–4728.e10. Available from: https://linkinghub.elsevier.com/retrieve/pii/S1097276524008633

39. Šviković S, Sale JE. The Effects of Replication Stress on S Phase Histone Management and Epigenetic Memory. J Mol Biol. 2017;429(13):2011–29.

40. Giunta S, Funabiki H. Integrity of the human centromere DNA repeats is protected by CENP-A, CENP-C, and CENP-T. Proc Natl Acad Sci U S A. 2017;114(8):1928–33.

41. Baranovskiy AG, Babayeva ND, Suwa Y, Gu J, Pavlov YI, Tahirov TH. Structural basis for inhibition of DNA replication by aphidicolin. Nucleic Acids Res. 2014;42(22):14013–21.

42. Vesela E, Chroma K, Turi Z, Mistrik M. Common chemical inductors of replication stress: Focus on cell-based studies. Biomolecules. 2017;7(1).

43. Ercilla A, Feu S, Aranda S, Llopis A, Brynjólfsdóttir SH, Sørensen CS, et al. Acute hydroxyurea-induced replication blockade results in replisome components disengagement from nascent DNA without causing fork collapse. Cellular and Molecular Life Sciences [Internet]. 2020;77(4):735–49. Available from: 10.1007/s00018-019-03206-1

44. Petermann E, Orta ML, Issaeva N, Schultz N, Helleday T. Hydroxyurea-Stalled Replication Forks Become Progressively Inactivated and Require Two Different RAD51-Mediated Pathways for Restart and Repair. Mol Cell. 2010;37(4):492– 502.

45. Lobbia VR, Trueba Sanchez MC, van Ingen H. Beyond the Nucleosome: Nucleosome-Protein Interactions and Higher Order Chromatin Structure. Vol. 433, Journal of Molecular Biology. Academic Press; 2021.

46. Musselman CA, Kutateladze TG. Characterization of functional disordered regions within chromatin-associated proteins. iScience [Internet]. 2021;24(2):102070. Available from: 10.1016/j.isci.2021.102070

47. Cho KF, Branon TC, Udeshi ND, Myers SA, Carr SA, Ting AY. Proximity labeling in mammalian cells with TurboID and split-TurboID. Nat Protoc [Internet]. 2020;15(12):3971–99. Available from: 10.1038/s41596-020-0399-0

48. Branon TC, Bosch JA, Sanchez AD, Udeshi ND, Svinkina T, Carr SA, et al. Efficient proximity labeling in living cells and organisms with TurboID. Nat Biotechnol. 2018;36(9):880–98.

49. Pan D, Walstein K, Take A, Bier D, Kaiser N, Musacchio A. Mechanism of centromere recruitment of the CENP-A chaperone HJURP and its implications for centromere licensing. Nat Commun [Internet]. 2019;10(1):1–18. Available from: 10.1038/s41467-019-12019-6

50. Khurana S, Varma D, Foltz DR. Contribution of CENP-F to FOXM1-Mediated Discordant Centromere and Kinetochore Transcriptional Regulation. Mol Cell Biol. 2024;44(6):209–25.

51. Zhou X, Wang R, Fan L, Li Y, Ma L, Yang Z, et al. Mitosin/CENP-F as a negative regulator of activating transcription factor-4. Journal of Biological Chemistry. 2005 Apr 8;280(14):13973–7.

52. Aytes A, Mitrofanova A, Lefebvre C, Alvarez MJ, Castillo-Martin M, Zheng T, et al. Cross-Species Regulatory Network Analysis Identifies a Synergistic Interaction between FOXM1 and CENPF that Drives Prostate Cancer Malignancy. Cancer Cell. 2014 May 12;25(5):638–51.

53. Nye J, Sturgill D, Athwal R, Dalal Y. HJURP antagonizes CENP-A mislocalization driven by the H3.3 chaperones HIRA and DAXX. PLoS One. 2018;13(10):1–21.

54. Lacoste N, Woolfe A, Tachiwana H, Garea AV, Barth T, Cantaloube S, et al. Mislocalization of the Centromeric Histone Variant CenH3/CENP-A in Human Cells Depends on the Chaperone DAXX. Mol Cell. 2014;53(4):631–44.

55. Shrestha RL, Ahn GS, Staples MI, Sathyan KM, Karpova TS, Foltz DR, et al. Mislocalization of centromeric histone H3 variant CENP-A contributes to chromosomal instability (CIN) in human cells. Oncotarget. 2017;8(29):46781–800.

56. Batenburg NL, Walker JR, Coulombe Y, Sherker A, Masson JY, Zhu XD. CSB interacts with BRCA1 in late S/G2 to promote MRN-A nd CtIP-mediated DNA end resection. Nucleic Acids Res. 2019 Nov 18;47(20):10678–92.

57. Batenburg NL, Walker JR, Noordermeer SM, Moatti N, Durocher D, Zhu XD. ATM and CDK2 control chromatin remodeler CSB to inhibit RIF1 in DSB repair pathway choice. Nat Commun [Internet]. 2017;8(1). Available from: 10.1038/s41467-017-02114-x

58. Boetefuer EL, Lake RJ, Fan HY. Mechanistic insights into the regulation of transcription and transcription-coupled DNA repair by Cockayne syndrome protein B. Vol. 46, Nucleic Acids Research. Oxford University Press; 2018. p. 7471–9.

59. Naughton C, Huidobro C, Catacchio CR, Buckle A, Grimes GR, Nozawa RS, et al. Human centromere repositioning activates transcription and opens chromatin fibre structure. Nat Commun [Internet]. 2022 Sep 24;13(1):5609. Available from: https://www.biorxiv.org/content/10.1101/2021.08.01.454615v2%0A https://www.biorxiv.org/content/10.1101/2021.08.01.454615v2.abstract

60. Ishikura S, Nakabayashi K, Nagai M, Tsunoda T, Shirasawa S. ZFAT binds to centromeres to control noncoding RNA transcription through the KAT2B-H4K8ac-BRD4 axis. Nucleic Acids Res. 2020;48(19):10848–66.

61. Shiromoto Y, Sakurai M, Minakuchi M, Ariyoshi K, Nishikura K. ADAR1 RNA editing enzyme regulates R-loop formation and genome stability at telomeres in cancer cells. Nat Commun. 2021 Dec 1;12(1).

62. Loyola A, Tagami H, Bonaldi T, Roche D, Quivy JP, Imhof A, et al. The HP1α-CAF1-SetDB1-containing complex provides H3K9me1 for Suv39-mediated K9me3 in pericentric heterochromatin. EMBO Rep. 2009;10(7):769–75.

63. Otake K, Ohzeki JI, Shono N, Kugou K, Okazaki K, Nagase T, et al. CENP-B creates alternative epigenetic chromatin states permissive for CENP-A or heterochromatin assembly. J Cell Sci. 2020;133(15):1–17.

64. Bailey AO, Panchenko T, Shabanowitz J, Lehman SM, Bai DL, Hunt DF, et al. Identification of the Post-translational Modifications Present in Centromeric Chromatin. Molecular & Cellular Proteomics [Internet]. 2016 Mar;15(3):918–31. Available from: https://linkinghub.elsevier.com/retrieve/pii/S1535947620336513

65. Sullivan BA, Karpen GH. Centromeric chromatin exhibits a histone modification pattern that is distinct from both euchromatin and heterochromatin. Nat Struct Mol Biol. 2004;11(11):1076–83.

66. Coker H, Brockdorff N. SMCHD1 accumulates at DNA damage sites and facilitates the repair of DNA double-strand breaks. J Cell Sci. 2014;127(9):1869– 74.

67. Tian T, Bu M, Chen X, Ding L, Yang Y, Han J, et al. The ZATT-TOP2A-PICH Axis Drives Extensive Replication Fork Reversal to Promote Genome Stability. Mol Cell. 2021 Jan 7;81(1):198–211.e6.

68. Challa K, Schmid CD, Kitagawa S, Cheblal A, Iesmantavicius V, Seeber A, et al. Damage-induced chromatome dynamics link Ubiquitin ligase and proteasome recruitment to histone loss and efficient DNA repair. Mol Cell. 2021;81(4):811–829.e6.

69. Hauer MH, Seeber A, Singh V, Thierry R, Sack R, Amitai A, et al. Histone degradation in response to DNA damage enhances chromatin dynamics and recombination rates. Nat Struct Mol Biol. 2017;24(2):99–107.

70. Singh RK, Gonzalez M, Kabbaj MHM, Gunjan A. Novel E3 ubiquitin ligases that regulate histone protein levels in the budding yeast saccharomyces cerevisiae. PLoS One. 2012 May 3;7(5).

71. Singh I, Ozturk N, Cordero J, Mehta A, Hasan D, Cosentino C, et al. High mobility group protein-mediated transcription requires DNA damage marker γ-H2AX. Cell Res. 2015 Jul 4;25(7):837–50.

72. Truch J, Downes DJ, Scott C, Gür ER, Telenius JM, Repapi E, et al. The chromatin remodeller ATRX facilitates diverse nuclear processes, in a stochastic manner, in both heterochromatin and euchromatin. Nat Commun. 2022 Dec 1;13(1).

73. Voon HPJ, Wong LH. New players in heterochromatin silencing: Histone variant H3.3 and the ATRX/DAXX chaperone. Nucleic Acids Res. 2015;44(4):1496–501.

74. Ratnakumar K, Duarte LF, LeRoy G, Hasson D, Smeets D, Vardabasso C, et al. ATRX-mediated chromatin association of histone variant macroH2A1 regulates α-globin expression. Genes Dev. 2012;26(5):433–8.

75. Ren W, Medeiros N, Warneford-Thomson R, Wulfridge P, Yan Q, Bian J, et al. Disruption of ATRX-RNA interactions uncovers roles in ATRX localization and PRC2 function. Nat Commun [Internet]. 2020;11(1):1–15. Available from: 10.1038/s41467-020-15902-9

76. Park J, Lee H, Han N, Kwak S, Lee HT, Kim JH, et al. Long non-coding RNA ChRO1 facilitates ATRX/DAXX-dependent H3.3 deposition for transcription-associated heterochromatin reorganization. Nucleic Acids Res. 2018;46(22):11759–75.

77. Huh MS, Ivanochko D, Hashem LE, Curtin M, Delorme M, Goodall E, et al. Stalled replication forks within heterochromatin require ATRX for protection. Cell Death Dis. 2016;7(5):1–12.

78. Raghunandan M, Yeo JE, Walter R, Saito K, Harvey AJ, Ittershagen S, et al. Functional cross talk between the Fanconi anemia and ATRX/DAXX histone chaperone pathways promotes replication fork recovery. Hum Mol Genet. 2021;29(7):1083–95.

79. Leung JWC, Ghosal G, Wang W, Shen X, Wang J, Li L, et al. Alpha thalassemia/mental retardation syndrome X-linked gene product ATRX is required for proper replication restart and cellular resistance to replication stress. Journal of Biological Chemistry [Internet]. 2013;288(9):6342–50. Available from: 10.1074/jbc.M112.411603

80. Juhász S, Elbakry A, Mathes A, Löbrich M. ATRX Promotes DNA Repair Synthesis and Sister Chromatid Exchange during Homologous Recombination. Mol Cell. 2018;71(1):11–24.e7.

81. Gulve N, Su C, Deng Z, Soldan SS, Vladimirova O, Wickramasinghe J, et al. DAXX-ATRX regulation of p53 chromatin binding and DNA damage response. Nat Commun. 2022;13(1):1–14.

82. Pladevall-Morera D, Munk S, Ingham A, Garribba L, Albers E, Liu Y, et al. Proteomic characterization of chromosomal common fragile site (CFS)-associated proteins uncovers ATRX as a regulator of CFS stability. Nucleic Acids Res. 2019;47(15):8004–18.

83. Nan X, Hou J, Maclean A, Nasir J, Jose Lafuente M, Shu X, et al. Interaction between chromatin proteins MECP2 and ATRX is disrupted by mutations that cause inherited mental retardation [Internet]. 2007. Available from: www.pnas.org/cgi/content/full/

84. Scott WA, Dhanji EZ, Dyakov BJA, Dreseris ES, Asa JS, Grange LJ, et al. ATRX proximal protein associations boast roles beyond histone deposition. 2021;1–30. Available from: 10.1371/journal.pgen.1009909

85. Ortega-Alarcon D, Claveria-Gimeno R, Vega S, Kalani L, Jorge-Torres OC, Esteller M, et al. Extending MeCP2 interactome: canonical nucleosomal histones interact with MeCP2. Nucleic Acids Res. 2024 Apr 24;52(7):3636–53.

86. Boetefuer EL, Lake RJ, Fan HY. Mechanistic insights into the regulation of transcription and transcription-coupled DNA repair by Cockayne syndrome protein B. Vol. 46, Nucleic Acids Research. Oxford University Press; 2018. p. 7471–9.

87. Cho I, Tsai PF, Lake RJ, Basheer A, Fan HY. ATP-Dependent Chromatin Remodeling by Cockayne Syndrome Protein B and NAP1-Like Histone Chaperones Is Required for Efficient Transcription-Coupled DNA Repair. PLoS Genet. 2013;9(4).

88. Batenburg NL, Mersaoui SY, Walker JR, Coulombe Y, Hammond-Martel I, Wurtele H, et al. Cockayne syndrome group B protein regulates fork restart, fork progression and MRE11-dependent fork degradation in BRCA1/2-deficient cells. Nucleic Acids Res. 2021;49(22):12836–54.

89. Batenburg NL, Thompson EL, Hendrickson EA, Zhu X. Cockayne syndrome group B protein regulates DNA double-strand break repair and checkpoint activation. EMBO J. 2015;34(10):1399–416.

90. Carraro M, Hendriks IA, Hammond CM, Solis-Mezarino V, Völker-Albert M, Elsborg JD, et al. DAXX adds a de novo H3.3K9me3 deposition pathway to the histone chaperone network. Mol Cell. 2023;83(7):1075–1092.e9.

91. Lewis PW, Elsaesser SJ, Noh KM, Stadler SC, Allis CD. Daxx is an H3.3-specific histone chaperone and cooperates with ATRX in replication-independent chromatin assembly at telomeres. Proc Natl Acad Sci U S A. 2010;107(32):14075–80.

92. Suzuki N, Nakano M, Nozaki N, Egashira SI, Okazaki T, Masumoto H. CENP-B Interacts with CENP-C Domains Containing Mif2 Regions Responsible for Centromere Localization. Journal of Biological Chemistry. 2004 Feb 13;279(7):5934–46.

93. Morozov VM, Giovinazzi S, Ishov AM. CENP-B protects centromere chromatin integrity by facilitating histone deposition via the H3.3-specific chaperone Daxx. Epigenetics Chromatin [Internet]. 2017;10(1):1–18. Available from: 10.1186/s13072-017-0164-y

94. Nechemia-Arbely Y, Miga KH, Shoshani O, Aslanian A, McMahon MA, Lee AY, et al. DNA replication acts as an error correction mechanism to maintain centromere identity by restricting CENP-A to centromeres. Nat Cell Biol [Internet]. 2019;21(6):743–54. Available from: 10.1038/s41556-019-0331-4

95. Hara M, Fukagawa T. Centromere maintenance during DNA replication. Nat Cell Biol. 2019;21(6):669–71.

96. Nathanailidou P, Dhakshnamoorthy J, Xiao H, Zofall M, Holla S, O’Neill M, et al. Specialized replication of heterochromatin domains ensures self-templated chromatin assembly and epigenetic inheritance. Proc Natl Acad Sci U S A. 2024 Feb 6;121(6).

97. Kim SM, Dubey DD, Huberman JA. Early-replicating heterochromatin. Genes Dev. 2003 Feb 1;17(3):330–5.

98. Gaggioli V, Lo CSY, Reverón-Gómez N, Jasencakova Z, Domenech H, Nguyen H, et al. Dynamic de novo heterochromatin assembly and disassembly at replication forks ensures fork stability. Nat Cell Biol. 2023;

99. Nikolov I, Taddei A. Linking replication stress with heterochromatin formation. Chromosoma [Internet]. 2016;125(3):523–33. Available from: 10.1007/s00412-015-0545-6

100. Bodor DL, Valente LP, Mata JF, Black BE, Jansen LET. Assembly in G1 phase and long-term stability are unique intrinsic features of CENP-A nucleosomes. Mol Biol Cell. 2013;24(7):923–32.

101. Stankovic A, Guo LY, Mata JF, Bodor DL, Cao XJ, Bailey AO, et al. A Dual Inhibitory Mechanism Sufficient to Maintain Cell-Cycle-Restricted CENP-A Assembly. Mol Cell. 2017 Jan 19;65(2):231–46.

102. Byun TS, Pacek M, Yee MC, Walter JC, Cimprich KA. Functional uncoupling of MCM helicase and DNA polymerase activities activates the ATR-dependent checkpoint. Genes Dev. 2005;19(9):1040–52.

103. Chang DJ, Lupardus PJ, Cimprich KA. Monoubiquitination of proliferating cell nuclear antigen induced by stalled replication requires uncoupling of DNA polymerase and mini-chromosome maintenance helicase activities. Journal of Biological Chemistry. 2006 Oct 27;281(43):32081–8.

104. Xu X, Duan S, Hua X, Li Z, He R, Zhaang Z. Stable inheritance of H3.3-containing nucleosomes during mitotic cell divisions. Nat Commun [Internet]. 2022;13(1):2514. Available from: http://www.ncbi.nlm.nih.gov/pubmed/35523900 %0A http://www.pubmedcentral.nih.gov/articlerender.fcgi?artid=PMC9076889

105. Morozov VM, Gavrilova E V., Ogryzko V V., Ishov AM. Dualistic function of Daxx at centromeric and pericentromeric heterochromatin in normal and stress conditions. Nucleus (United States). 2012;3(3).

106. Leen Lam A, Boivin CD, Bonney CF, Katharine Rudd M, Sullivan BA, Henikoff S. Human centromeric chromatin is a dynamic chromosomal domain that can spread over noncentromeric DNA [Internet]. 2006. Available from: www.pnas.orgcgidoi10.1073pnas.0507947103

107. Black EM, Giunta S. Repetitive fragile sites: Centromere satellite DNA as a source of genome instability in human diseases. Genes (Basel). 2018;9(12).

108. Bodor DL, Rodríguez MG, Moreno N, Jansen LET. Analysis of protein turnover by quantitative SNAP-based pulse-chase imaging. Curr Protoc Cell Biol. 2012;(SUPPL.55):1–42.

109. Zhou Y, Zhou B, Pache L, Chang M, Khodabakhshi AH, Tanaseichuk O, et al. Metascape provides a biologist-oriented resource for the analysis of systems-level datasets. Nat Commun [Internet]. 2019;10(1). Available from: 10.1038/s41467-019-09234-6

110. Shrestha RL, Balachandra V, Kim JH, Rossi A, Vadlamani P, Sethi SC, et al. The histone H3/H4 chaperone CHAF1B prevents the mislocalization of CENP-A for chromosomal stability. J Cell Sci. 2023;136(10).

