## Supplementary material for "Replication stress alters CENP-A nucleosome stability during S phase": Lee_et_al_RS_CENPA_SUPP_INFO_initial_submission

### Table of Contents: Total 11 pages


- **SI Figure 1:** Flow cytometry analysis of cell cycle progression during replication stress induction.
- **SI Figure 2:** Validation of doxycycline-inducible expression of TurboID cell lines and enrichment of biotinylated interacting proteins.
- **SI Figure 3:** Proximity labeling by Chromatin-TurboID reveals distinct and specific histone H3 variant interactions and diverse functions.
- **SI Figure 4:** Proximity labeling by Chromatin-TurboID reveals global proteome changes in both centromeric and general chromatin under replication stress.
- **SI Figure 5:** DAXX is required for ATRX accumulation at centromeres.
- **SI Figure 6:** Validation of siRNA knockdowns of ATRX and DAXX.
- **SI Figure 7:** Replication stress leads to increased levels of centromeric H3K9me3.
- **SI Figure 8:** Cell cycle progression under siDAXX or ATRX depleted condition in subsequent G1 phase
- **SI Figure 9:** Validation of doxcycyline induction of CENP-A overexpression.

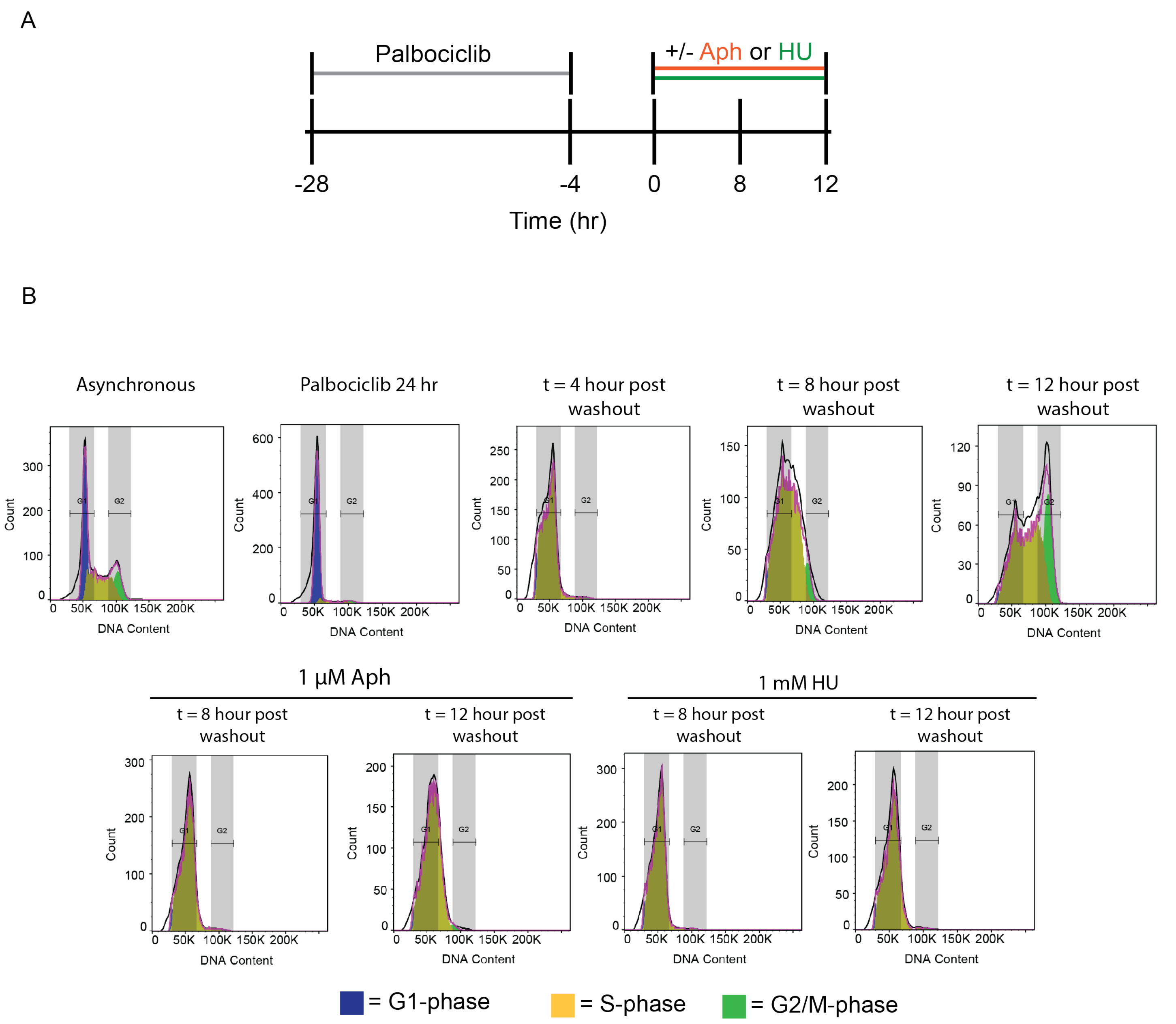


**SI Figure 1: Flow cytometry analysis of cell cycle progression during replication stress induction. A)** Schematic of experimental design. **B)** Representative FACs profiles of hTERT-RPE1 cells were treated with either 1 mM hydroxyurea (HU) or 1 μM Aphidicolin (Aph) for four hours at timepoint t = 4 hours post-washout. Cell cycle positions of cells were analyzed by DNA content by flow cytometry. Experiments were performed with three independent biological replicates.

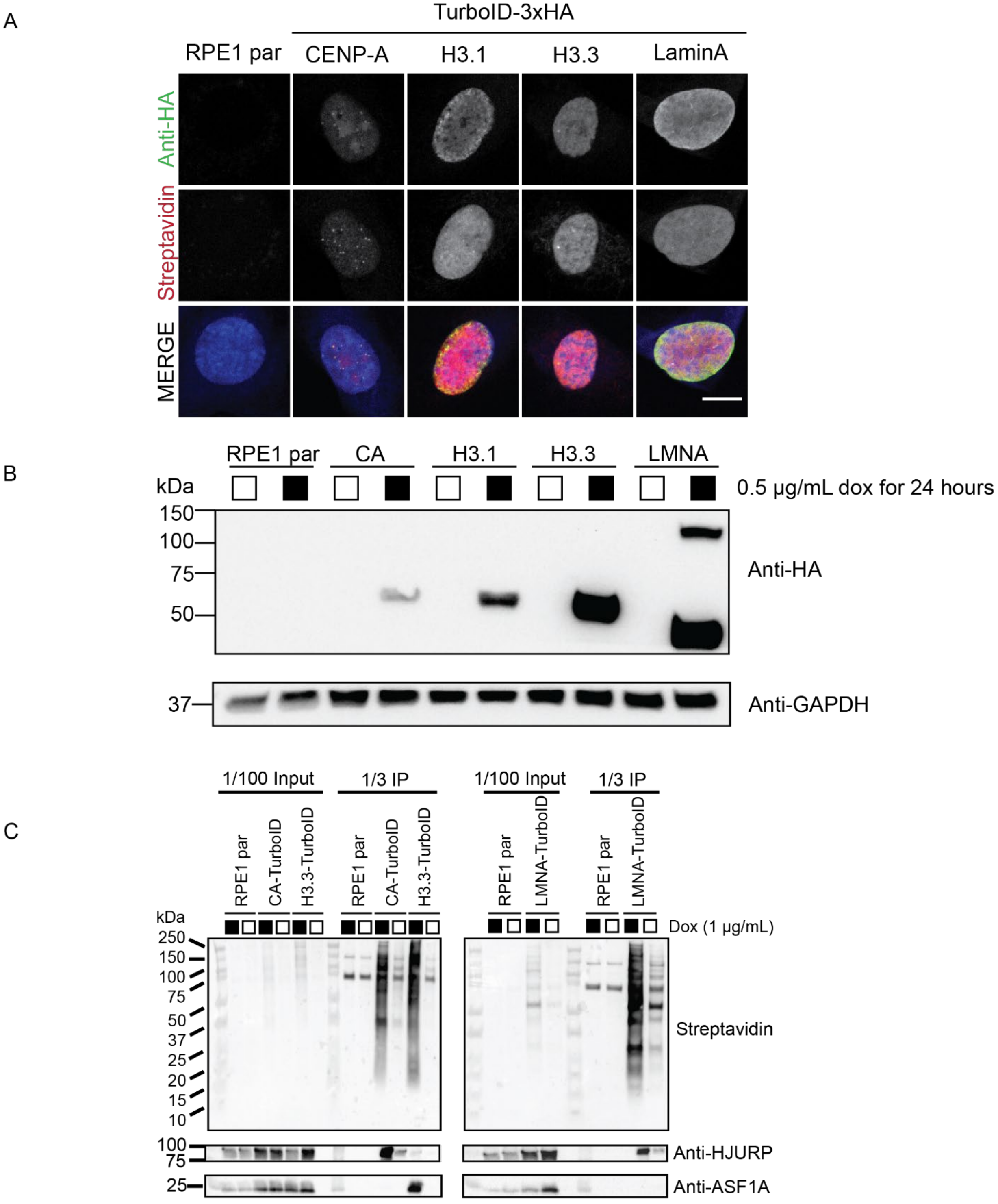


**SI Figure 2: Validation of doxycycline-inducible expression of TurboID cell lines and enrichment of biotinylated interacting proteins. A)** TurboID cell lines were treated with doxycycline (1 µg/mL) for 24 hours, and in vitro biotinylation labeling was conducted at 100 µM for 1 hour prior to indirect immunofluorescence staining (scale bar = 10 µm). **B)** Whole cell extracts of TurboID cell lines treated with doxycycline (0.5 µg/mL) for 24 hours, and *in vitro* biotinylation labeling was conducted at 100 µM for 1 hour. **C)** Doxycycline induction of Histone-TurboID fusion cells and validation of pulldowns of known interaction partners. Doxycycline induction was conducted at 0.5 µg/mL for 24 hrs and treated with 1 mM biotin for 1 hr prior to cell harvest.

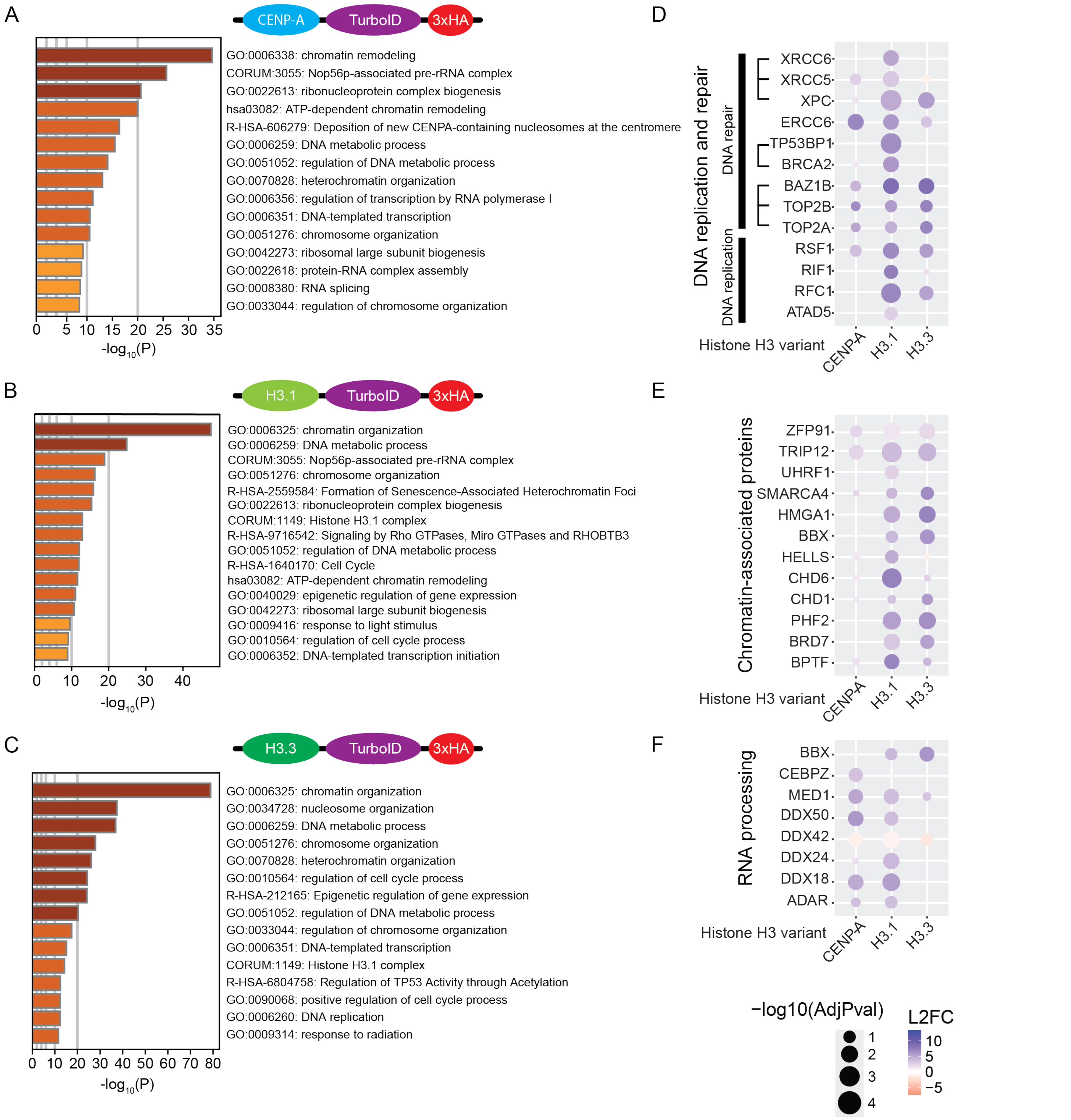


**SI Figure 3: Proximity labeling by Chromatin-TurboID reveals distinct and specific histone H3 variant interactions and diverse functions. A)** Gene Ontology (GO) enrichment analysis of histone H3 variant-enriched protein-protein interactions by AP-MS including **A)** CENP-A, **B)** H3.1, and **C)** H3.3. Strength is the -$Log_{10}$ (observed/expected) and is a measure of enrichment effect. **D)** Bubble plots of protein groups identified by AP-MS of randomly cycling histone-turboID constructs, and protein groups were grouped based on GO-term defined pathways including **D)** DNA replication and repair, **E)** Chromatin-association, and **F)** RNA processing. Brackets indicate known protein complexes. Three biological replicates were analyzed with three technical replicates per biological replicate. $Log_2$ Fold Change (L2FC) is of [Histone/LaminA control]. Adjusted *p* values were calculated based on bonferroni correction.

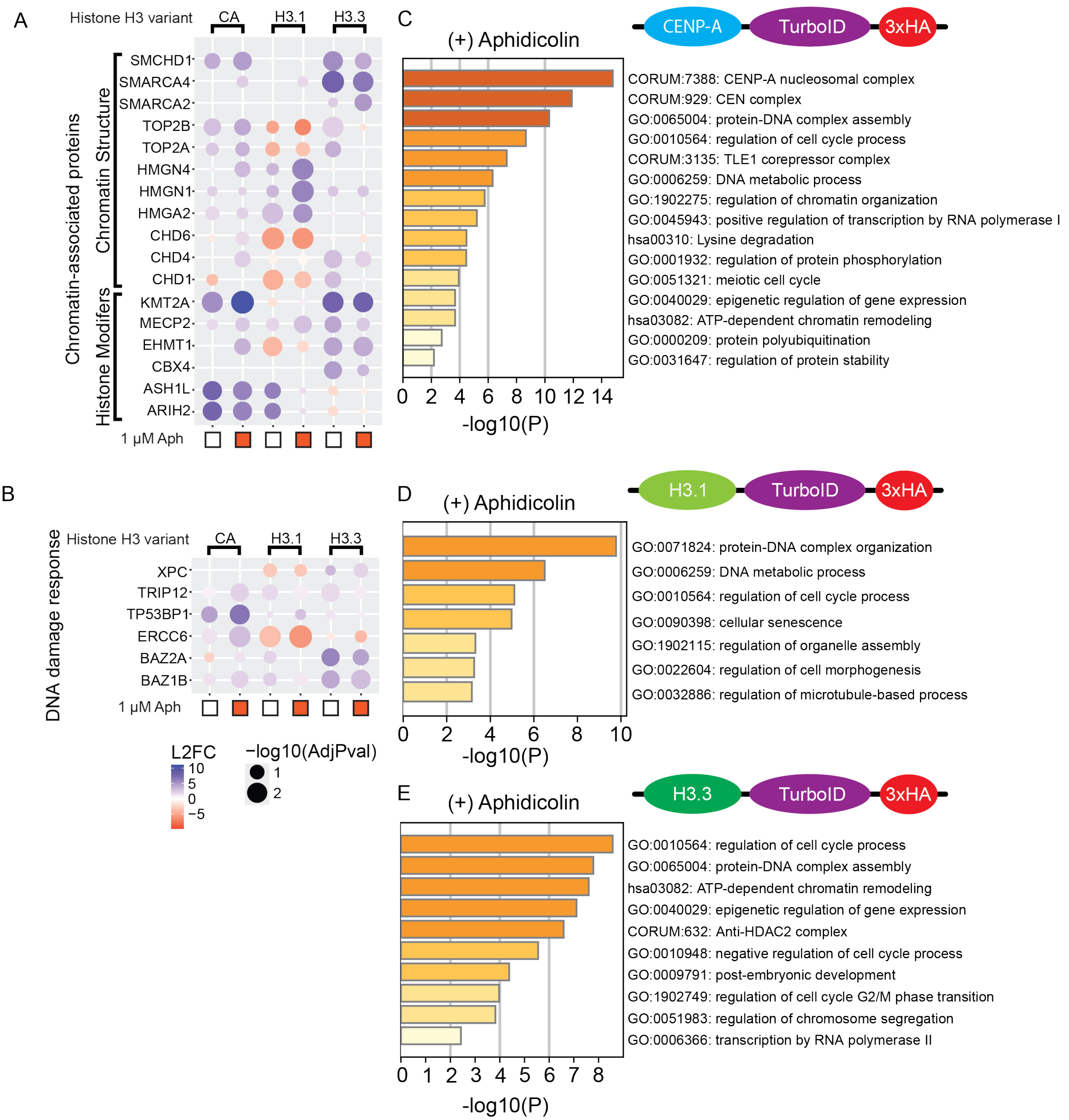


**SI Figure 4: Proximity labeling by Chromatin-TurboID reveals global proteome changes in both centromeric and general chromatin under replication stress. A)** Bubble plots of representing pulldowns of histone-dependent interactions by proximity labeling under normal replication and replication stress identified by BP-MS using histone-turboID constructs. Protein groups were grouped based on GO-term defined pathways including **A)** Chromatin-association and **B)** DNA damage response. Three biological replicates were analyzed with three technical replicates per biological replicate. $Log_2$ Fold Change (L2FC) is of [Histone/LaminA control]. Adjusted *p* values were calculated based on bonferroni correction. **C)** Gene Ontology (GO) enrichment analysis of histone H3 variant-enriched protein-protein interactions including **C)** CENP-A, **D)** H3.1, and **E)** H3.3 under aphidicolin-induced replication stress. Strength is the $-Log_{10}$ (observed/expected) and is a measure of enrichment effect.

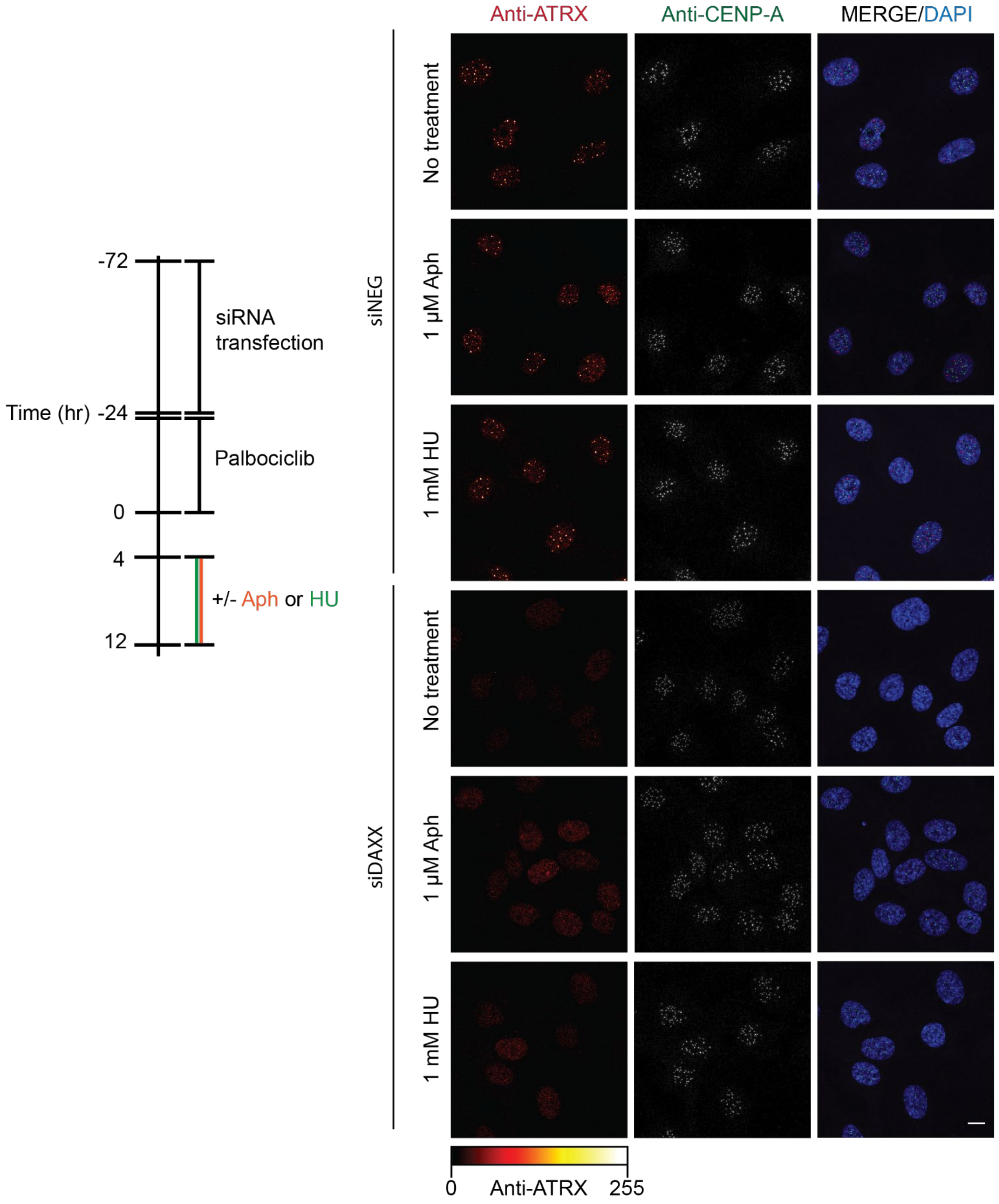


**SI Figure 5: DAXX is required for ATRX accumulation at centromeres.** Schematic of experimental design (left) and representative images of hTERT-RPE1 cells transfected with siNEG control or siDAXX under DNA replication or Aph/HU-induced replication stress (right). Cells were co-stained for CENP-A and ATRX. Scale bar is 10 μm.

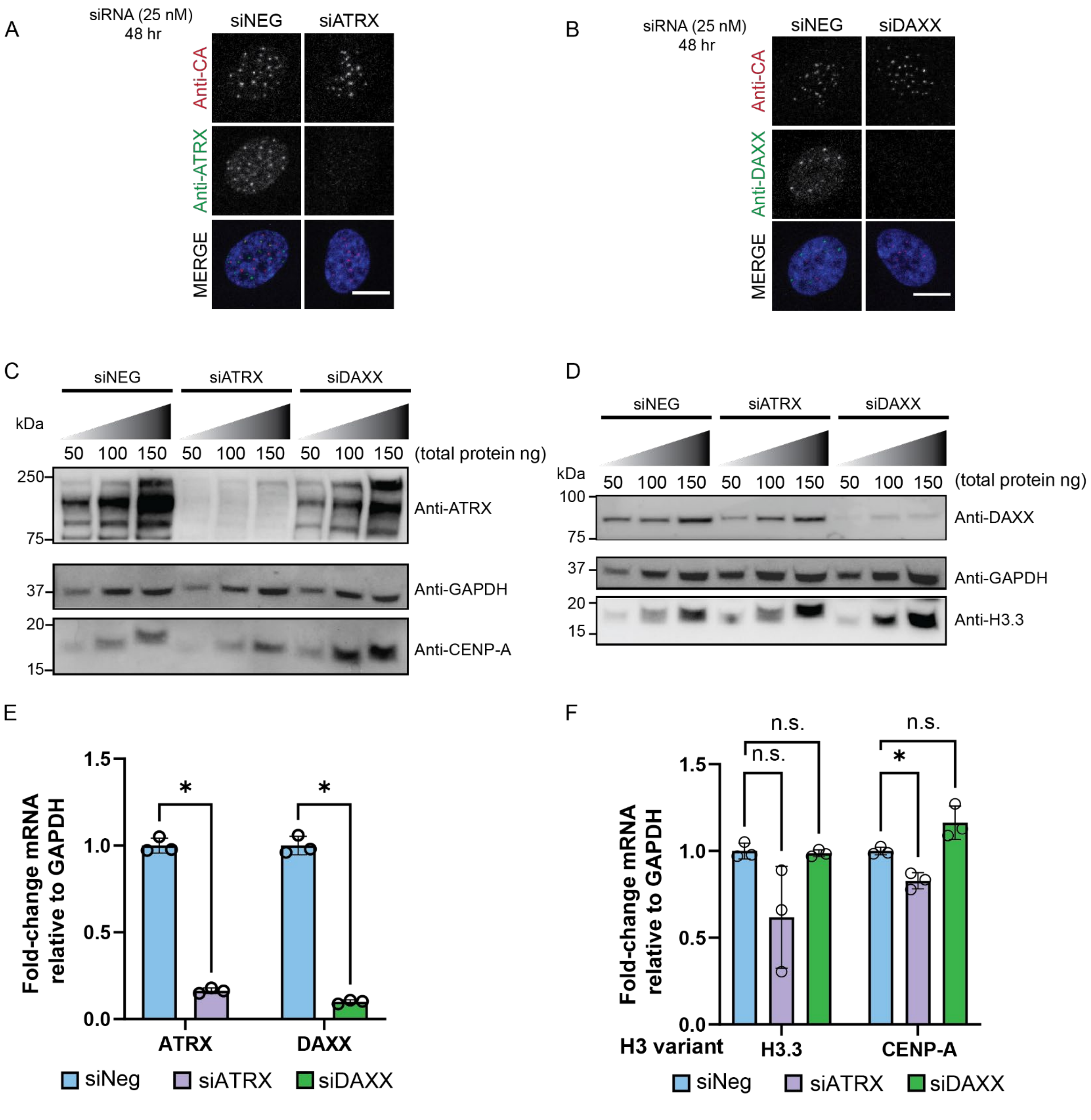


**SI Figure 6: Validation of siRNA knockdown of ATRX and DAXX. A)** Representative images of siATRX knockdown in RPE1 cells co-stained with ATRX and CENP-A. **B)** Representative images of siDAXX knockdown in RPE1 cells co-stained with DAXX and CENP-A. **C)** Total protein levels of CENP-A from siRNA depletion of ATRX and DAXX in RPE1 cells. **D)** Total protein levels of H3.3 from siRNA depletion of ATRX and DAXX in RPE1 cells. **E)** qPCR quantification of ATRX or DAXX expression relative to GAPDH under siATRX or DAXX depleted conditions. **F)** qPCR quantification of histone H3 variant H3.3 or CENP-A expression relative to GAPDH under siATRX or DAXX depleted conditions. Not significant (n.s.) = $p > 0.05$, $p \leq 0.05$ = *, $p \leq 0.01$ = **, $p \leq 0.001$ = ***. qPCR experiments were conducted with three independent biological replicates. Scale bar is 10 µm. Error bars represent standard deviation from the mean.

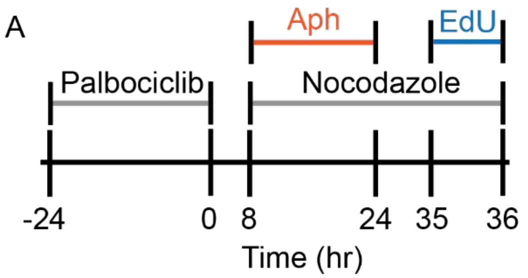


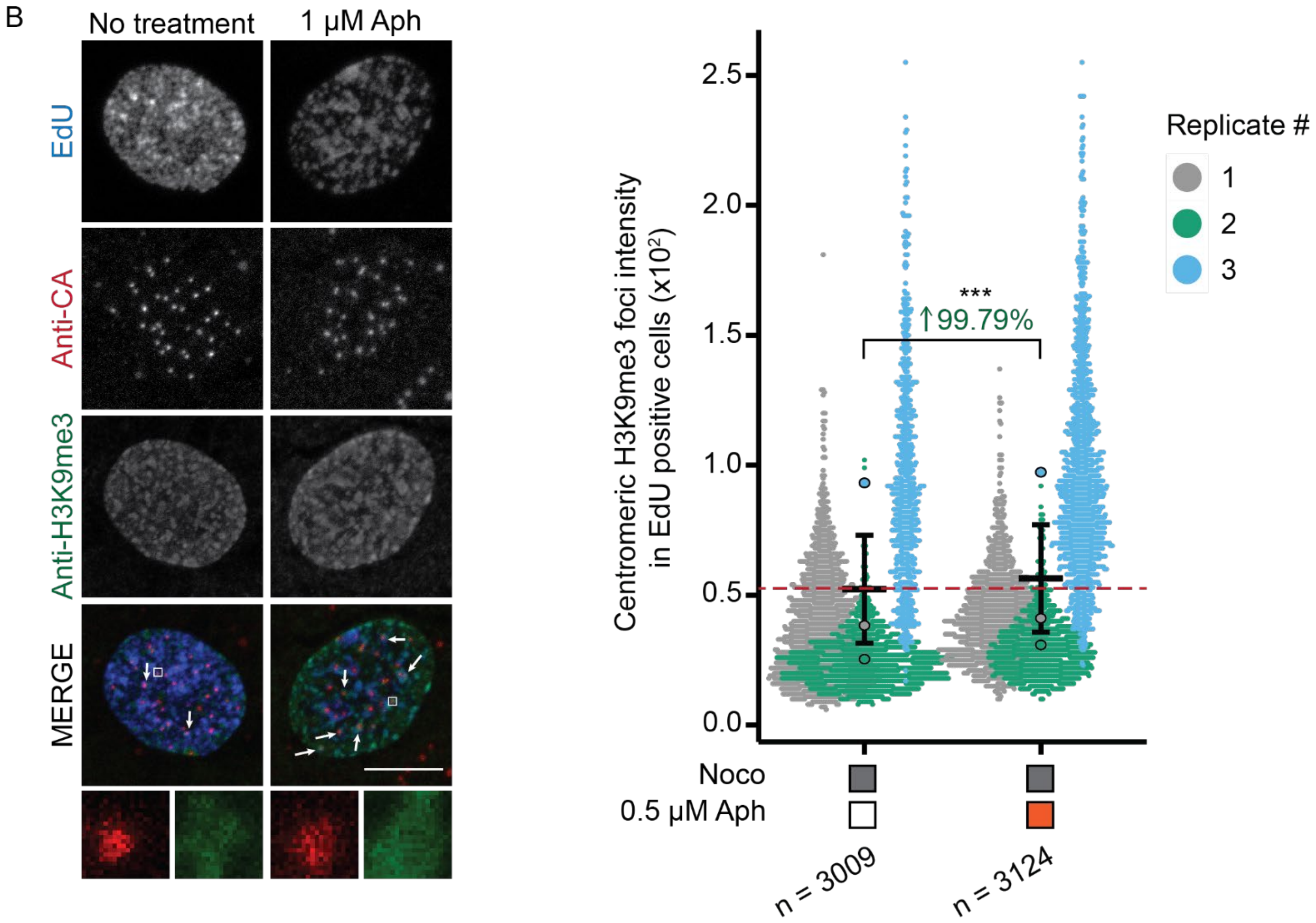


**SI Figure 7: Replication stress leads to increased levels of centromeric H3K9me3. A)** Schematic of experimental design. **B)** Representative images of RPE1 cells under replication or Aph-induced stress conditions (left) and quantification of centromeric H3K9me3 levels using CENP-A as a centromere reference marker (right). Dashed line indicates baseline average intensity relative to nocodazole only control. Experiments were performed three independent biological replicates and n = represents number of CENP-A foci measured. Not significant (n.s.) = $p > 0.05$, $p \leq 0.05$ = *, $p \leq 0.01$ = **, $p \leq 0.001$ = ***. Scale bar = 10 µm.

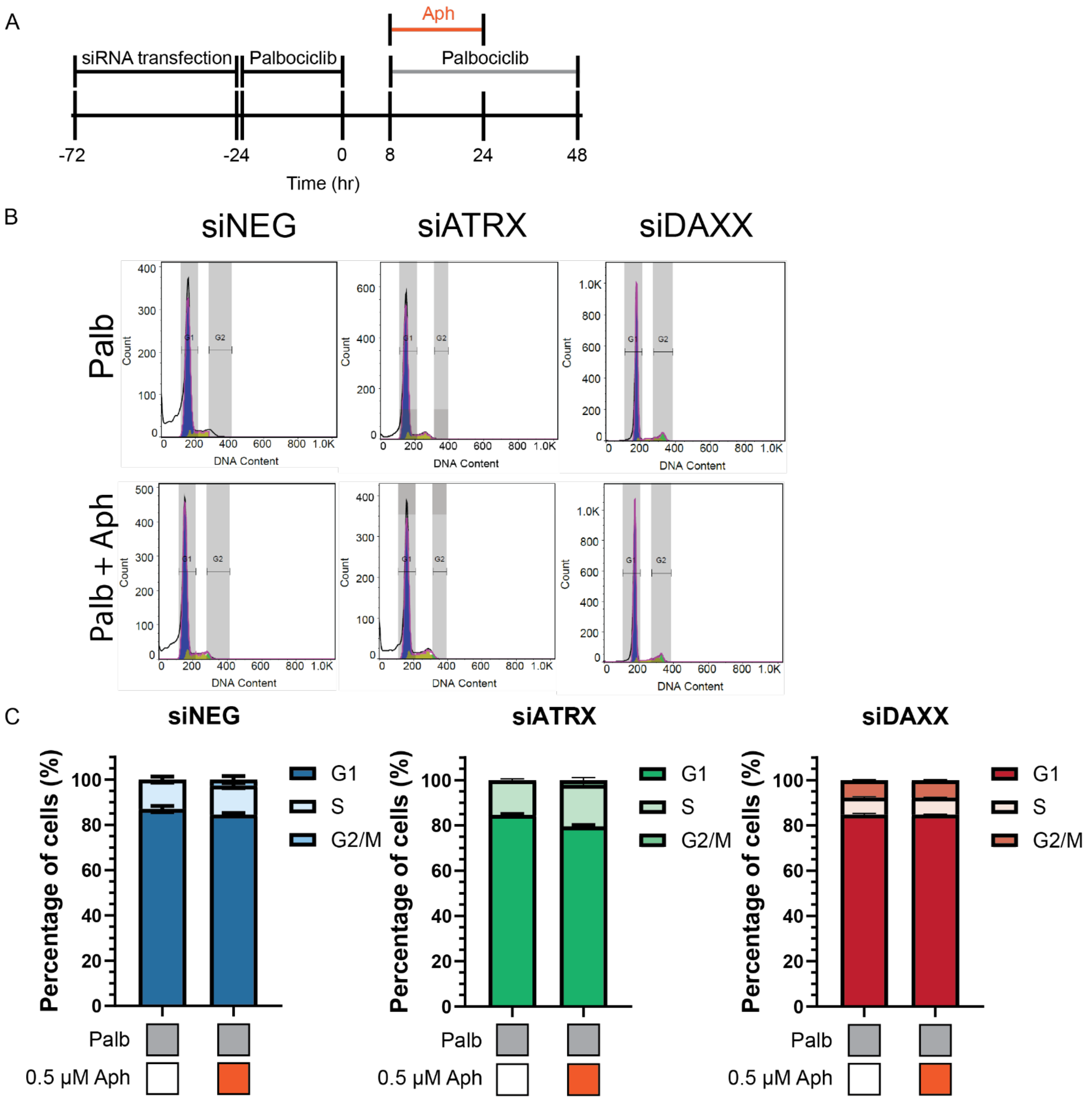


**SI Figure 8: Cell cycle progression under siDAXX or ATRX depleted condition in subsequent G1 phase. A)** Schematic of experimental design. **B)** Representative flow cytometry profiles by DNA content. Purple represents G1-phase population, yellow represents S-phase population, and green represents G2/M phase population. **C)** Quantification of cell population in cell cycle phase under siNEG control (left), siATRX (middle), or siDAXX (right) depleted conditions in RPE1 cells arrested. Cell cycle phases were analyzed by flow cytometry based on DNA content. Experiments were performed with three independent biological replicates. Error bars represent standard error of the mean.

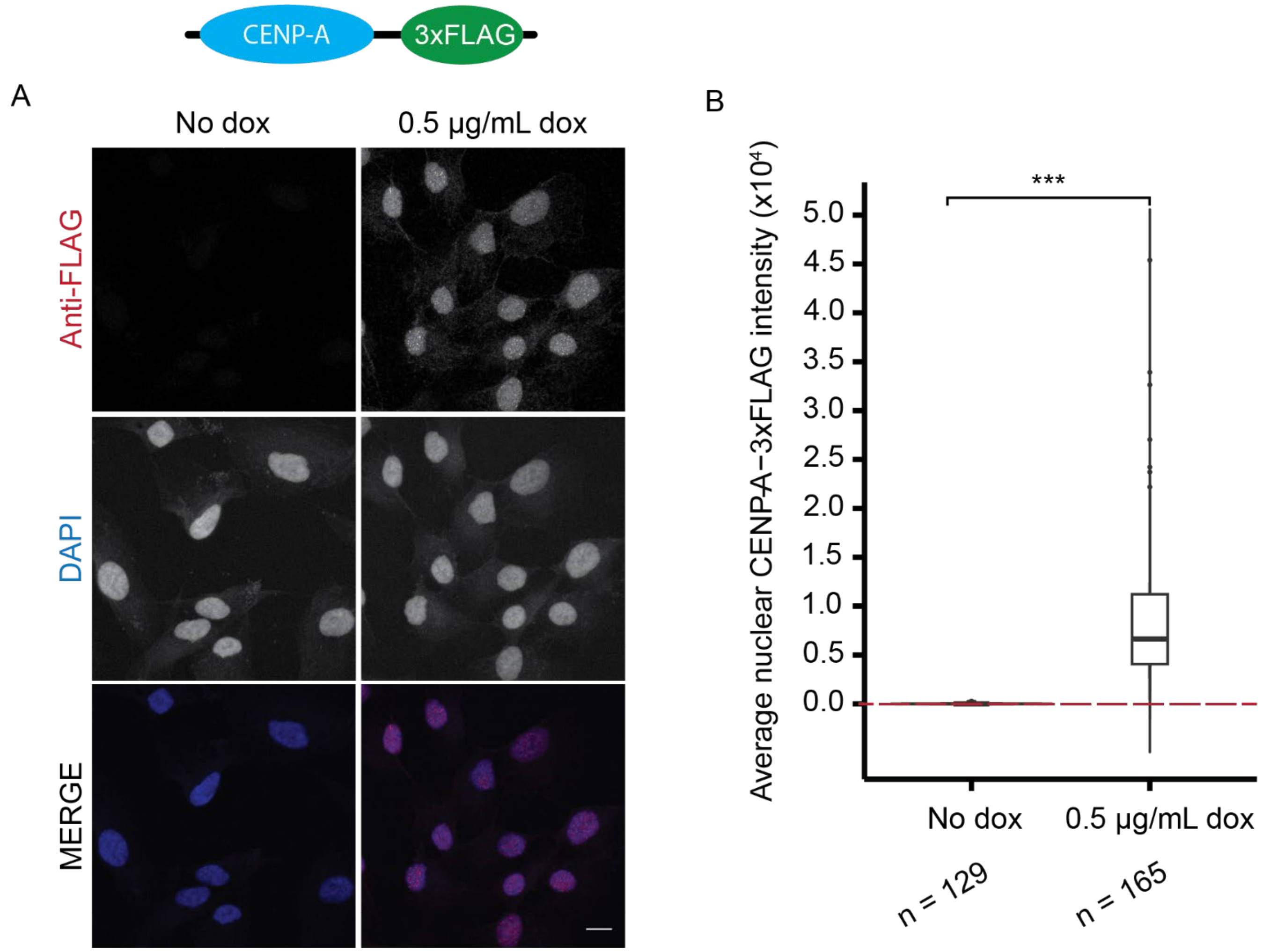


**SI Figure 9: Validation of doxcycyline induction of CENP-A overexpression. A)** Representative images of RPE1 cells were stably expressing CENP-A-3xFLAG under a doxcycline-inducible promoter. Stable cell lines were treated with 0.5 µg/mL doxycycline (dox) for 24 hours. Cells were immunofluorescent stained against FLAG-epitope to validate overexpression of CENP-A. Scale bar = 10 µm. **B)** Quantification of nuclear CENP-A-FLAG signal in response to dox induction. Dashed line indicates baseline average intensitiy relative to no dox control. n = represents number of cells measured per group. Not significant (n.s.) = $p > 0.05$, $p \leq 0.05$ = *, $p \leq 0.01$ = **, $p \leq 0.001$ = ***. Experiments were performed with three independent biological replicates. Less than 1% of cells exhibited leaky expression of CENP-A-3xFLAG in no dox control.
